# The recreational-to-habitual shift in psychostimulant use is an economic demand parameter that is unrelated to drug consumption levels (under normal and punishment conditions)

**DOI:** 10.64898/2026.05.19.726350

**Authors:** Martin O. Job, Indu Mithra Madhuranthakam, Kona Basak, Shakil Ahmed, Azim Uddin, Mst Afroza Alam Tumpa, Alida Jimenez, Rachel Cherry, Adriana Rodriguez, Maria Chowdhury, Thomas M. Keck

**Author notes:** Corresponding author: (MOJ).

## Abstract

**Rationale:** The progression of psychostimulant abuse is associated with a shift from recreational to habitual use (R2H-shift). Because this R2H-shift can be modeled using behavioral economics, we developed a novel Behavioral Economic model for the Analysis of Self-administration Time-curve (BEAST) to obtain R2H-shift variable(s). The relationship(s) between R2H-shift variables and drug intake (under normal and/or punishment conditions) is/are unknown. Our goal was to determine if the R2H-shift variable and intake variables obtained during the initial self-administration training phase were related to 1) drug intake at that time, and subsequent drug intake under 2) normal, 3) punishment, 4) post-punishment, and 5) price-constrained conditions.

**Method:** Long Evans rats self-administered methamphetamine (METH, males n = 16, females n = 14), sucrose (males n = 22, females n = 22) and/or saline (males n = 3, females n = 10) under FR1 for 6 h per day for 20 days to obtain 1) followed by the assessment of subsequent drug intake under different conditions (2-5 above). We obtained all variables referenced above. We determined the relationships between all variables (multivariate analysis).

**Results:** There were no sex differences detected in the METH and sucrose studies. For METH and sucrose, prior drug intake levels could predict drug intake under normal/punishment but not under price-constrained conditions. The R2H-shift variable could predict drug intake under a consumption-price curve but could not predict intake under normal/punishment conditions.

**Conclusions:** While related to economic demand, the recreational-to-habitual shift rate was unrelated to drug intake levels (under normal and punishment conditions).

## Introduction

Drug use progression is thought to be associated with a shift from recreational to habitual use (R2H-shift) which is sometimes described also as goal-directed to compulsive use. This shift is thought to involve progressive recruitment of dorsal striatal regions in drug use (Everitt and Robbins, 2013, 2005; Lüscher et al., 2020; Zapata et al., 2010). However, this R2H-shift is not well characterized. Understanding this will be imperative if we are to understand the mechanisms governing the progression of drug use and the typology of drug users. This knowledge, in turn, may be beneficial for individualized or personalized pharmacotherapeutic intervention strategies to address the drug use epidemic.

To characterize this R2H-shift, we carefully examined what occurs in drug self-administration. In drug self-administration, a subject performs a task (such as an active lever press) to obtain a reward (such as a drug infusion). Over time, the subject repeats this behavior and gains more experience with the drug. When a task is repeated over time, a shift to *automaticity* occurs (Haith and Krakauer, 2018) – an R2H-shift occurs. Automaticity changes behavior in at least two ways including 1) permitting appropriate actions to be selected with less cognitive load/effort (less cognitive price) than before, and 2) allowing habit formation. This reduction in cognitive *price* is related to a corresponding shift from prefrontal cortex to basal ganglia regulation of behavior (Ashby et al., 2010; Barnett et al., 2023; Everitt and Robbins, 2005; Isoda and Hikosaka, 2011; Lipton et al., 2019; Meyer et al., 2016; Morris et al., 2017) (Figure 1A-B) and is associated with an R2H-shift. This reduction in cognitive price (and hence this R2H-shift) is a behavioral economic phenomenon. Because the self-administration time curve is a type of behavioral economic system, R2H-shift may be captured using behavioral economic analysis of the drug self-administration time curve.

**Figure 1.**
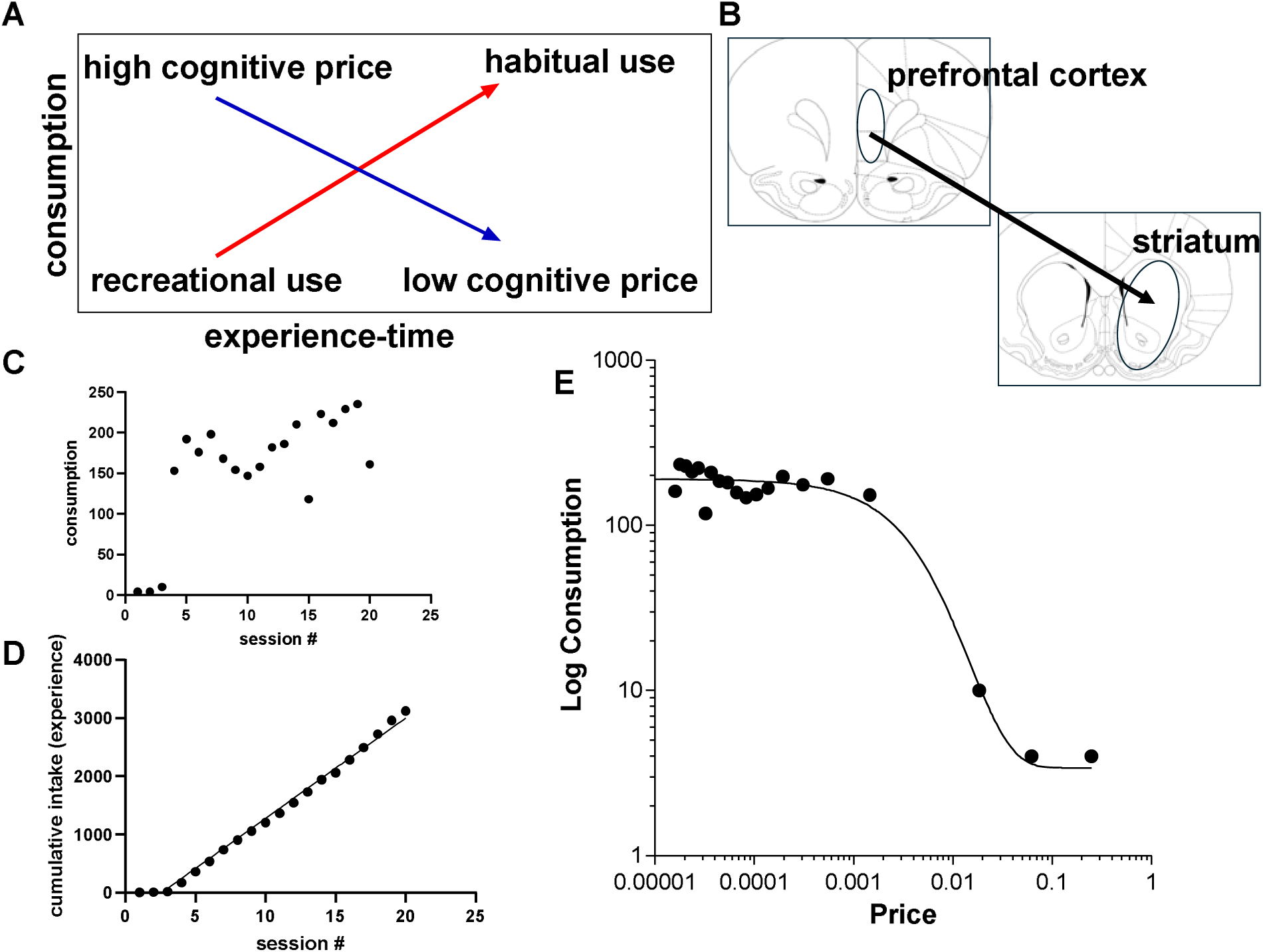
The new Behavioral Economic model for the Analysis of Self-administration Time-curve (BEAST). Graph A highlights the phenomenon of automaticity – the subject repeats a task to obtain a reinforcer over time, gaining experience simultaneously, with the ultimate development of automaticity which includes habit formation and a decrease in cognitive price (cognitive effort). Graph B highlights evidence in the literature that suggests that this decrease in cognitive price represents a shift from prefrontal cortex activity (associated with high energy expenditure/high price) to striatum (associated with lower energy/low price) to minimize the price required to execute behavior. This decrease in cognitive price associated with shifts from recreational-to-habitual drug use is a behavioral economic activity. C, D and E are data from the same individual showing how we developed BEAST. C is a plot of time on the x-axis and consumption on the y-axis. D is a plot of time on the x-axis and cumulative intake (experience) on the y-axis. With increases in experience-time, (decreases in cognitive price), the price required for behavioral activity also decreases. Thus, time and experience (experience-time) may be conceptualized as price constructs. Because the price required to complete these tasks decreases as experience-time increases, we expressed price as the reciprocal value of time (time includes # of sessions and hours per session) multiplied by the reciprocal of the value of experience. We expressed experience as cumulative activity from the beginning of the self-administration time curves. We plotted log consumption on the y-axis and fitted the resulting curve using the exponential model – this yielded a demand curve (E) from which we were able to obtain behavioral economic parameters Q_0_ (consumption at zero price), alpha (demand elasticity) and k (a scaling constant reflecting the consumption range). From alpha and k, we obtained the essential value (eValue). Unlike demand elasticity, which is a change in consumption when price increases, the R2H-shift rate/price represent a change in consumption from recreational to habitual that occurs when price decreases. We derived R2H-shift rate/price are from the natural log of the inverse of estimated eValue and Pmax.

However, there are no behavioral economic analytical models for the drug self-administration time course and so using experience-time as a price construct (Figure 1C-D), we developed a new model (Figure 1E). This novel model, termed Behavioral Economic Analysis of Self-Administration Time curve (BEAST, Figure 1E) utilizes the same data already employed in current model but it also accounts for a price construct that includes both time and experience. From this new BEAST model we can simultaneously estimate Q_0_ (drug consumption at zero price), essential value (eValue, demand elasticity) and Pmax (maximum price where demand shifts from inelastic to elastic). But because the price (experience-time) is decreasing, not increasing, over time, we can quantify the direction of the variables that have to do with slope (rate of change) and price, expressing them as 1/eValue and 1/Pmax. These variables should capture the R2H-shift.

The role of the R2H-shift in predicting behaviors after the initial drug self-administration time curve is unknown. Our goal was to determine if R2H-shift variable(s) obtained during self-administration training phase was/were related to 1) drug intake at that time, and subsequent drug intake under 2) normal, 3) punishment, 4) post-punishment, and 5) price-constrained conditions. For this study, we allowed different groups of male and female Long Evans rats to self-administer methamphetamine (METH) or sucrose (natural reinforcer) and saline. We assessed the self-administration time curve using current and BEAST. Our methods, results, discussion and conclusions are below.

## Methods

### Animals

The Rowan University Animal Care and Use Committee approved all procedures and treatments. All procedures and treatments followed the guidelines outlined in the National Institutes of Health (NIH) *Guide for the Care and Use of Laboratory Animals*. The animals, male and female Long-Evans rats were obtained from Charles River Laboratories (Wilmington, MA). All animals were adults and age-matched (56–62 days old at the time of arrival). Rats were housed in pairs under a 12-hour light/dark cycle with ad libitum access to food and water.

### Animal use

We employed a total of eighty-seven (87) rats spread across three major self-administration experiments, namely METH (N = 30), sucrose (N = 44) and saline (N = 13). The distribution by biological sex was as follows: METH (males n = 16, females n = 14), sucrose (males n = 22, females n = 22) and saline (males n = 3, females n = 10). For the METH (and saline) studies, we inserted a catheter into the rat’s external jugular vein. For jugular vein catheterization surgical procedures, see (Castaneda and Job, 2026; Job et al., 2020; Job and Katz, 2019). These rats were singly housed after the surgeries to preserve the catheters. There were no surgeries for the sucrose group. All animals were allowed to recover from surgery for approximately one-to-two weeks before self-administration experiments. For self-administration procedure and equipment used, see (Castaneda and Job, 2026).

### Experimental Design

For the experimental design, see Figure 2. After the animals had recovered from the surgical procedure, they were allowed to self-administer 0.1 mg/kg/infusion of METH (or sucrose or saline) on a Fixed Ratio = 1 (FR1) schedule for 6 hours per day for 5 days a week (no self-administration experiments on the weekend) for 4 weeks (20 sessions) (training phase, Figure 2A). Afterwards, they were allowed to self-administer drugs in a multicomponent schedule (Figure 2B) which contained 11 blocks of 30 min sessions each separated by 3 min for a total of 330 min (6 hours). The total intake during these blocks was designated intake post-training (Figure 2B) to represent intake (under normal conditions/ no punishment) after the drug self-administration time course. Note that after the training phase the multicomponent block design was used for the remainder of the experiment. After this 5-day post-training multicomponent sessions/day (normal intake), the subject underwent a punishment regimen (in a single day, Figure 2C) with footshock intensities increasing over the 11-30 min blocks as follows: 0.0, 0.1, 0.2, 0.3, 0.4, 0.5, 0.6, 0.7, 0.8, 0.9 and 1.0 mA. We conducted the footshock regimen in a single day to align with typical demand curve analysis. Following the punishment regimen, subjects underwent a post-punishment session (Figure 2D) in which they were allowed to self-administer drugs in the same block design as above (11-30 min sessions) but they were not subjected to any punishment (in this phase, they were allowed to recover from the punishment phase) - we obtained the intake-post punishment variable. Finally, the subject’s demand curves were assessed (Figure 2E, in a single day to allow standardized comparison with the punishment regimen) with price = effort requirements (Fixed Ratio, FR) and with FR increasing over the 11-30 min blocks as follows: FR 1, 3, 5, 10, 20, 50, 100, 200, 300, 500 and 1000.

**Figure 2.**
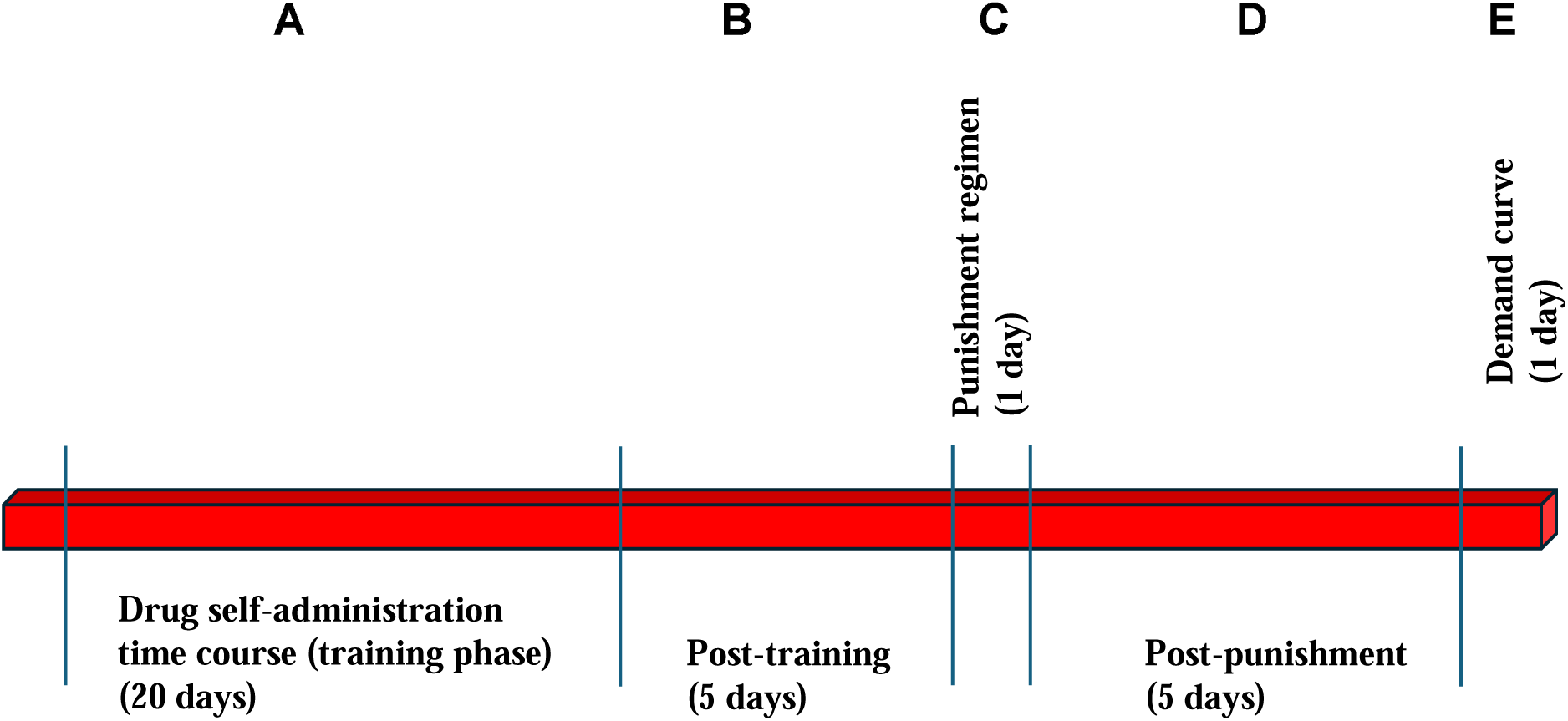
Experiment design: Long Evans rats self-administered methamphetamine (METH) or sucrose under FR1 (Fixed Ratio = 1 which means 1 active lever press to obtain 1 infusion of the reinforcer) for 6 h per day for 20 days – this is the drug self-administration training phase (A). Then subjects were allowed to self-administer drug on a multicomponent schedule (11 sub-sessions per day) for 5 days (post-training phase, B). Afterwards, subjects self-administered drug under a punishment phase (C) with increasing footshock intensities (mA) as follows: 0.0, 0.1, 0.2, 0.3, 0.4, 0.5, 0.6, 0.7, 0.8, 0.9 and 1.0 (11 footshock intensities over 11 components in a single day). After the footshock phase (1 day) subjects were allowed to self-administer drug as they did in the post-training phase (5 days, no punishment) –this is the post-punishment phase (D). After this phase, subjects self-administered drug under increases in price. The price included FR requirements to obtain the same reinforcer. These prices were as follows: FR 1, 3, 5, 10, 20, 50, 100, 200, 300, 500 and 1000 (11 price requirements over 11 components in a single day). From this design, we obtained ten (10) variables. Our goal was to determine the relationship between variables, especially to determine the correlates of the R2H-shift variables. R2H-shift stands for recreational-to-habitual shift (of drug use).

### Behavioral Economic Analysis of Self-Administration Time curve (BEAST)

For BEAST (Figure 1E), we conceptualized price as a construct that includes time itself and experience (cumulative/aggregate drug intake increasing with increments in time). Our rationale for this price construct is that with increases in time (session #) and increases in experience with the drug, the subject reduces the price of the drug to allow the R2H-shift (Figure 1). Thus, we conceptualized experience and time as price constructs using a formula as follows:

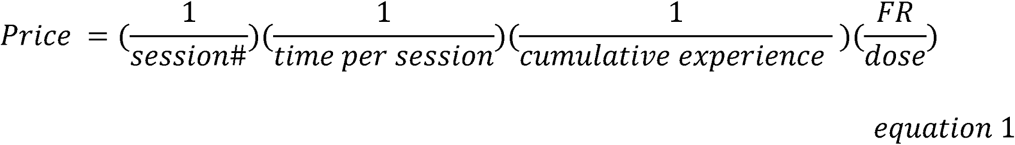

Because all subjects self-administered same dose of drugs (dose) for 6h per session (time per session) at the same FR, we derived price (for this specific study) as follows:

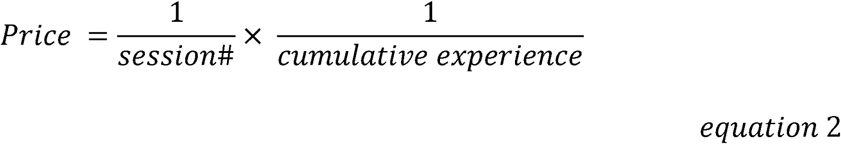

Note that for current demand curves used in the field, the Price (demand curves) = FR/dose. Equation 1 includes this current price construct but has also been expanded to include experience-time. The demand curve (Fig 1E) was fitted using the exponential function (Hursh and Silberberg, 2008) shown below:

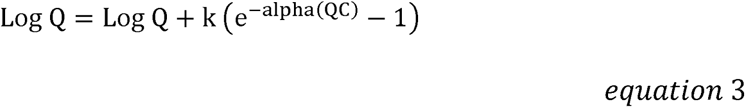

Where Q represents consumption of the reinforcer, C represents cost (price), Q_0_ represents consumption at zero (or no) cost, alpha represents demand elasticity and is a fitted parameter related to the decline in consumption with increased cost, and k is a scaling constant that corresponds to the consumption range.

The demand curve variable-alpha is inversely related to how much work the subject is willing to do to defend consumption when prices are increased. We calculated the essential value or eValue as follows:

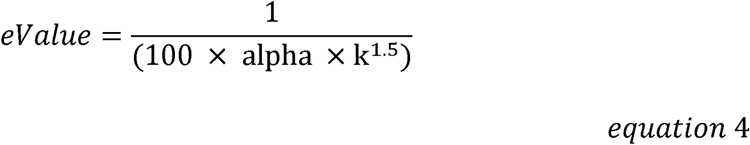

We calculated the Pmax (maximum price where demand shifts from inelastic to elastic) as follows:

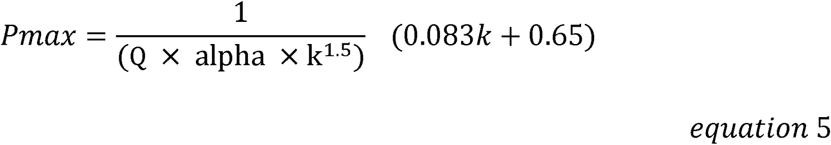

Because BEAST follows a reverse pattern relative to conventional demand curves, the R2H-shift rate was calculated as follows:

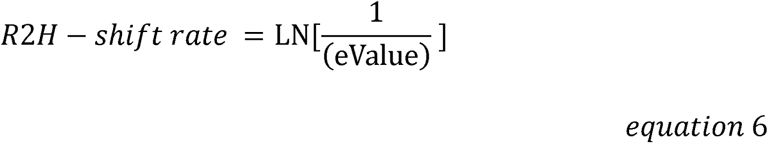

Where LN is natural logarithm. A higher R2H-shift rate implies a steeper slope for the change from recreational to habitual drug use (or steeper change in behavior in response to changes in drug price).

The R2H-shift price was calculated as follows:

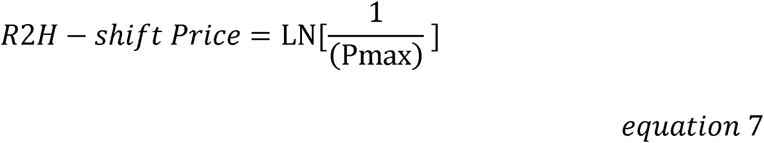

Where LN is natural logarithm. A higher R2H-shift price implies the higher the specific price at which the subject shifts from recreational to habitual drug use.

### Exclusion Criteria

For BEAST, we employed the exponential function (equation 3), noting the R^2^ for goodness of fit for all individuals (N = 87). We excluded subjects in which 1) the exponential model could not fit the demand curve, and/or 2) the alpha and/or eValue and/or Pmax were negative values.

### Variables

We obtained/derived 10 variables from several phases of the experiment design (Figure 2).

Training phase (Figure 2A): We obtained intake-training as the summation of intake during the 20 sessions of the training phase (cumulative intake). We obtained slope (sometimes termed escalation) as the rate of change of intake over the training phase time (using linear regression analysis). We obtained the mean of intake in the last 3 days of the training phase (MILD3)- this variable is used in many studies of this kind to establish stable intake before pharmacological/behavioral manipulation. For this phase, we also obtained Q_0_, R2H-shift rate and price (see calculations above).

Post-training (Figure 2B): We obtained intake post-training as the summation of intake during the 5 days post-training.

Punishment phase (Figure 2C): We obtained intake-punishment as the area under the consumption-punishment intensity curve (the summation of intake during the 1 day of the punishment regimen).

Post-punishment (Figure 2D): We obtained intake post-punishment as the summation of intake during the 5 days post-punishment.

Demand curve (Figure 2E): We obtained intake-PRICE as the area under the consumption-price curve (summation of intake during the 1 day of the demand curve analysis).

### Statistical analysis

GraphPad Prism v 10 (GraphPad Software, San Diego, CA), SigmaPlot 14.5 (Systat Software Inc., San Jose, CA) and JMP Pro v 18 (SAS Institute Inc., Cary, NC) were employed for statistical analysis. Data were expressed as mean ± SEM. We employed GraphPad templates for behavioral economic analysis/graphing from https://ibrinc.org/behavioral-economics-tools/. We used a calculator for eValue and Pmax which we downloaded from https://kuscholarworks.ku.edu/bitstreams/989121cf-1bd2-40d4-8205-7fdbc564d0df/download. We generated R2H-shift rate and R2H-shift price from eValue and Pmax (see equations 6-7).

### Assessment for Biological Sex differences

We employed a cluster-based approach to determine if there were biological sex differences across many different variables simultaneously in line with the Mapping of Intrinsic Sex Similarities as Integral qualities of Normalized Groups (MISSING) model (Job, 2024; Showell and Job, 2024; Tigano and Job, 2025, 2024; Yao and Job, 2024). For this, we employed principal component analysis (PCA) to reduce the above 10 variables into principal components (PCs) followed by gaussian mixtures model clustering of the PCs to determine if there were distinct clusters. If there are biological sex or individual differences, we should see clusters distinguished by sex or group. The criteria for cluster identification was that the clusters had to be clearly distinct and non-overlapping on a 3-dimensional space defined by principal components that together accounted for all the variability within the data set (PC 1, PC 2 and PC 3). For similar approaches, see (Ben-Hur and Guyon, 2003; Ruan et al., 2025).

### Multivariate analysis

To determine the relationships (if any) between variables, we conducted multivariate analysis on a total of 10 variables per subject: intake-training, slope, MILD3, Q_0_, R2H-shift rate, R2H-shift price, intake post-training, intake-punishment, intake post-punishment and intake-PRICE.

## Results

### Employment of BEAST to obtain R2H-shift variables for METH, sucrose and saline studies

The self-administration time demand curves for METH for males (n = 16) and females (n = 14) are shown in Figure 3 and Figure 4, respectively. The self-administration time demand curves for sucrose for males (n = 22) and females (n = 22) are shown in Figure 5 and Figure 6, respectively. For METH males, the exponential model (Hursh and Silberberg, 2008) for demand curves could either not fit or yielded negative values for eValue for the following individuals: rat2, rat3, rat4, rat9, rat10, rat11, rat12, rat13, rat15 and rat 16 (see graphs in Figure 3). For METH females, the exponential model could fit all demand curves (see Figure 4). For sucrose males, the exponential model for demand curves could either not fit or yielded negative values for eValue for the following individuals: rat33 and rat43 (see graphs in Figure 5). For sucrose males, the exponential model for demand curves could either not fit or yielded negative values for eValue for the following individuals: rat56 and rat72 (see graphs in Figure 6). For saline, the exponential model for demand curves could either not fit or yielded negative values for eValue for all subjects (Figure S1, Supplemental Figure 1), and all were excluded from further analysis. Thus the data analyzed for METH included n = 6 males and n = 14 females while the data analyzed for sucrose included n = 20 males and n = 20 females. For individuals included in the analysis, a summary of all variables obtained are shown in Table S1 for METH males and females, Table S2 for sucrose males and Table S3 for sucrose females.

**Figure 3.**
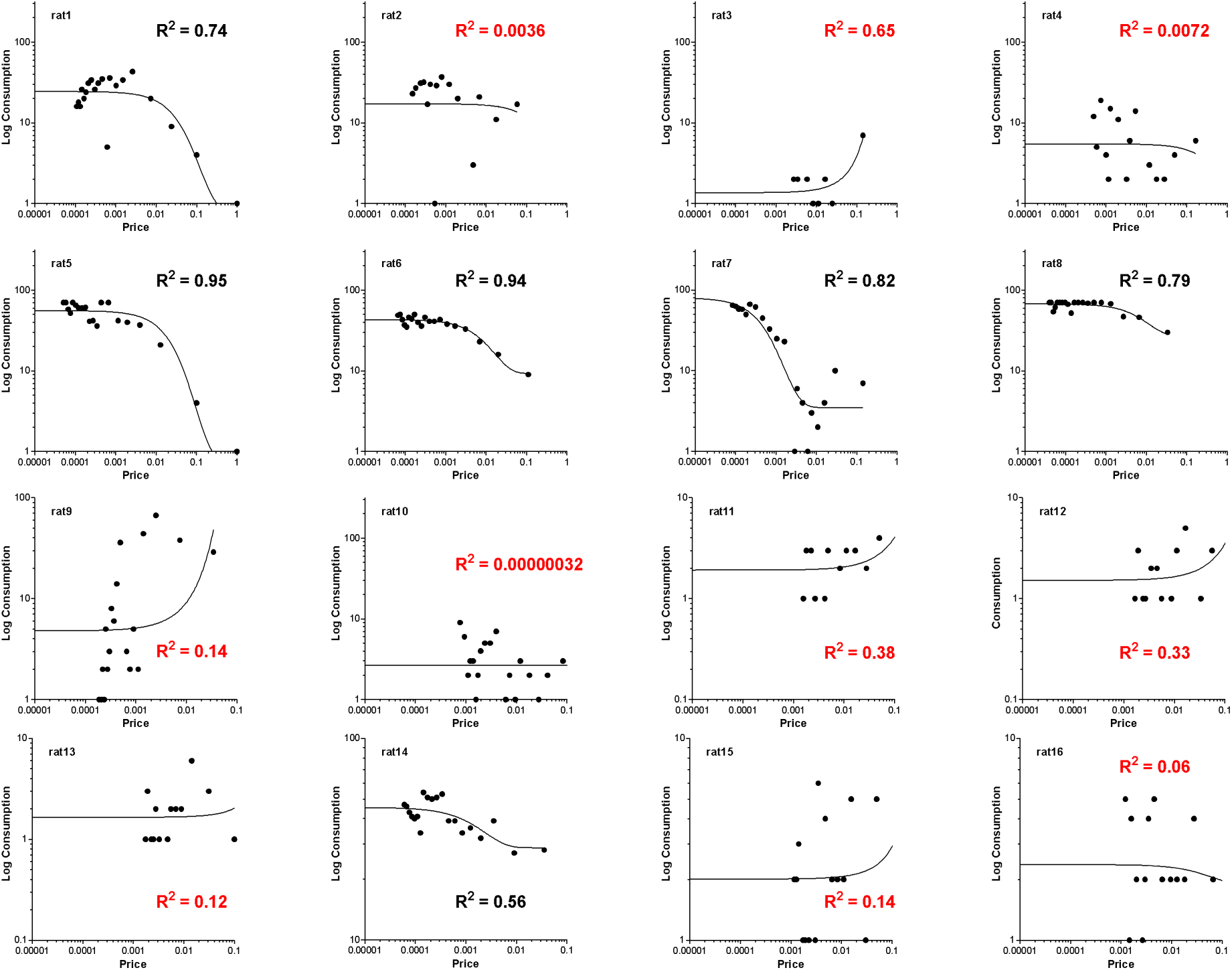
The demand curves of individuals (males) derived from METH self-administration time curve analysis using the new Behavioral Economic model for the Analysis of Self-administration Time-curve (BEAST). Each graph represents an individual. The n = 16 male rats that self-administered METH were labeled as rat1 – rat16. The R^2^ for goodness of fit is written in each graph. Note that the exponential model for demand curves could not fit and/or yielded negative values for eValue and Pmax for the following individuals: rat2, rat3, rat4, rat9, rat10, rat11, rat12, rat13, rat15 and rat 16 (R^2^ shown in red). These individuals were excluded from further analysis. The data for behavioral economic parameters obtained for included subjects are revealed in Table S1 (Supplemental Table 1).

**Figure 4.**
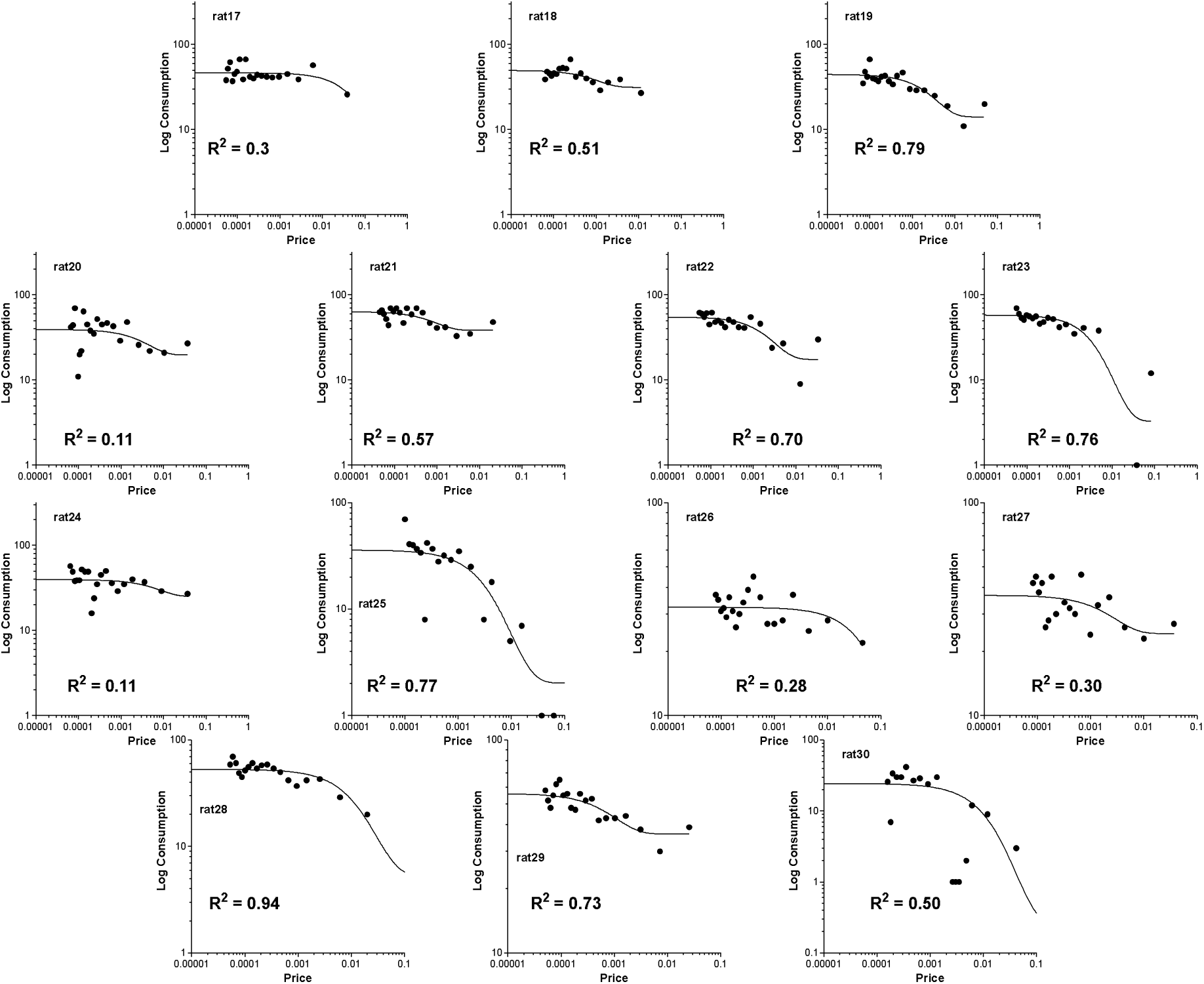
The demand curves of individuals (females) derived from METH self-administration time curve analysis using the new Behavioral Economic model for the Analysis of Self-administration Time-curve (BEAST). Each graph represents an individual. The n = 14 female rats that self-administered METH were labeled as rat17 – rat30. The R^2^ for goodness of fit is written in each graph. Note that the exponential model for demand curves could fit all subjects. The data for behavioral economic parameters obtained for all subjects are revealed in Table S1 (Supplemental Table 1).

**Figure 5.**
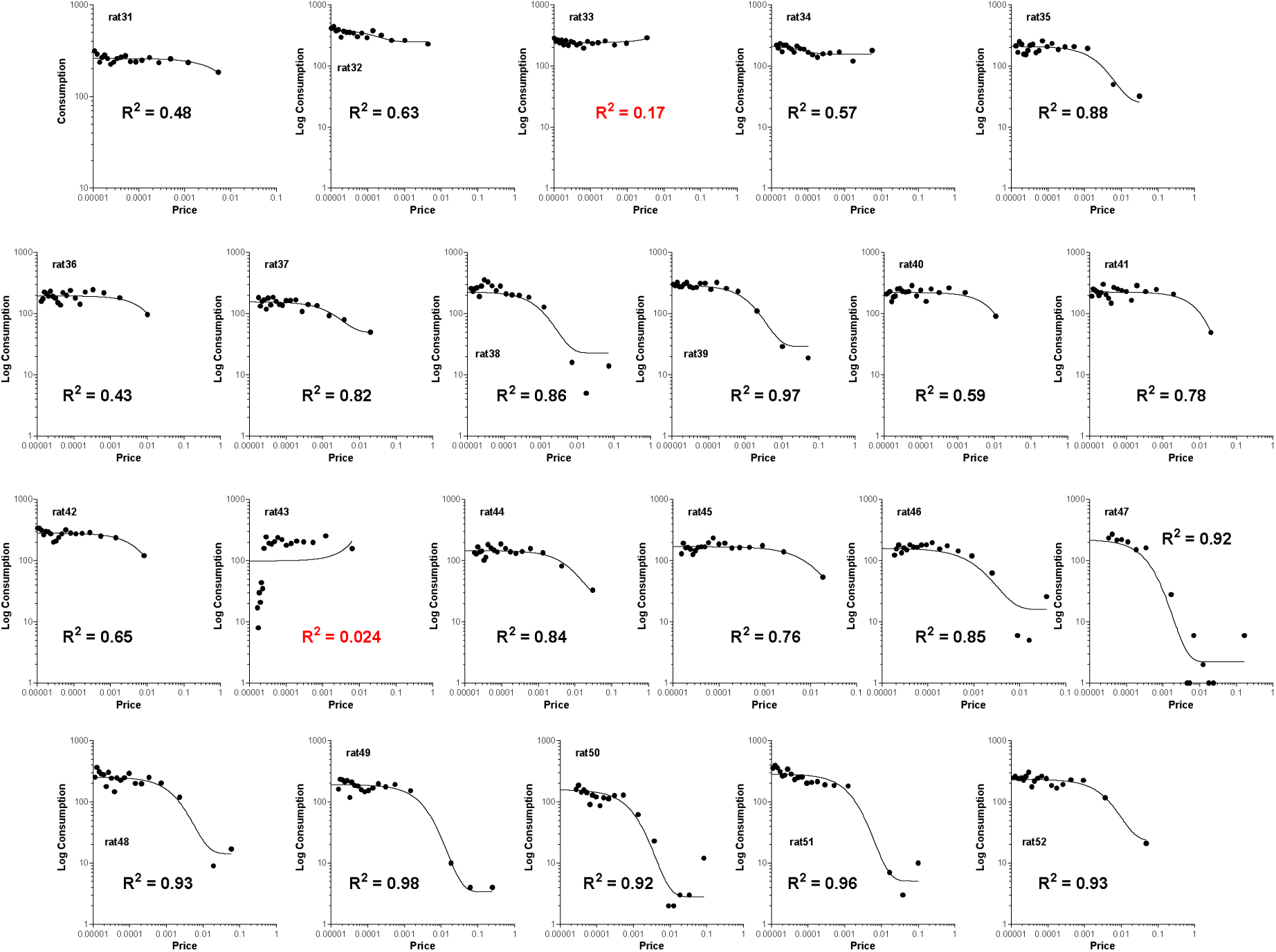
The demand curves of individuals (males) derived from sucrose self-administration time curve analysis using the new Behavioral Economic model for the Analysis of Self-administration Time-curve (BEAST). Each graph represents an individual. The n = 22 male rats that self-administered sucrose were labeled as rat31 – rat52. The R^2^ for goodness of fit is written in each graph. Note that the exponential model for demand curves could not fit and/or yielded negative values for eValue and Pmax for n = 2 individuals: rat33 and rat43 (R^2^ shown in red). These two individuals were excluded from further analysis. The data for behavioral economic parameters obtained for included subjects are revealed in Table 2 (Supplemental Table 2).

**Figure 6.**
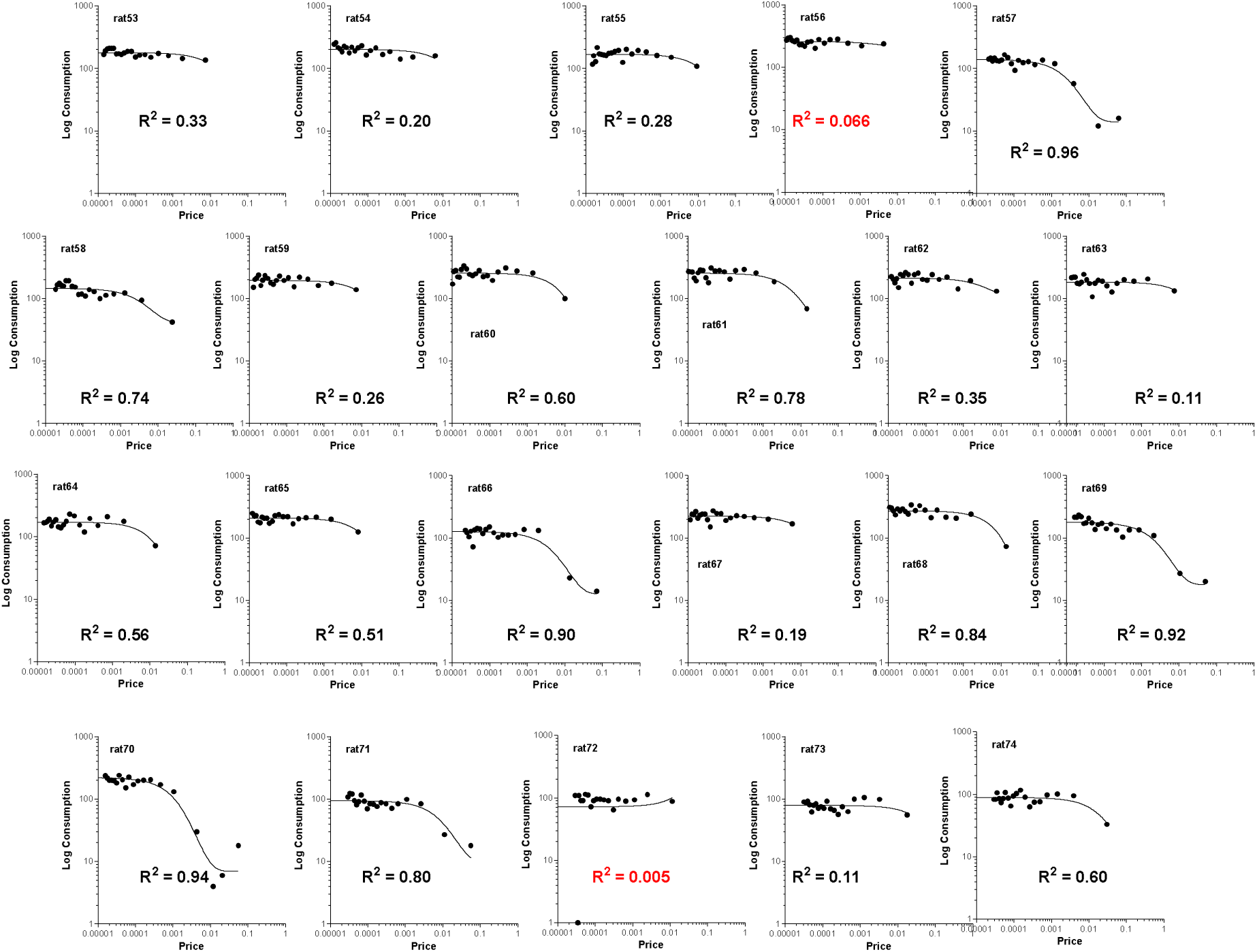
The demand curves of individuals (females) derived from sucrose self-administration time curve analysis using the new Behavioral Economic model for the Analysis of Self-administration Time-curve (BEAST). Each graph represents an individual. The n = 22 female rats that self-administered sucrose were labeled as rat53 – rat74. The R^2^ for goodness of fit is written in each graph. Note that the exponential model for demand curves could not fit and/or yielded negative values for eValue and Pmax for n = 2 individuals: rat56 and rat72 (R^2^ shown in red). These two individuals were excluded from further analysis. The data for behavioral economic parameters obtained for included subjects are revealed in Table 3 (Supplemental Table 3).

### Employment of PCA/ gaussian mixtures model clustering of PCs for the analysis of sex differences for METH and sucrose studies

Based on information that we have reported about the limitations of SABV (sex as a biological variable) revealed by the MISSING model (Job, 2024; Showell and Job, 2024; Tigano and Job, 2025, 2024; Yao and Job, 2024), we employed a cluster-based model (Figure 7), instead of current SABV (Figure S2-3), to simultaneously assess all 10 variables at the level of the individual first and at the level of biological sex second. Thus, we employed principal component analysis to reduce the 10 variables obtained into principal components (PCs). Afterwards, we conducted gaussian mixtures model clustering of the derived principal components to determine if we could detect distinct clusters of males and females. For the METH study, the proportion of variance for the PCs are shown in Figure 7A. Figure 7B shows the relative loading of the variables into PC 1 and PC 2. Gaussian mixtures model clustering revealed only one cluster (Figure 7C) – males and females were not clearly distinct. The cluster composition (males and females) relative to PC 1 and PC 2 are shown in Figure 7D. The cluster composition (males and females) relative to PC 1 and PC 2 and PC 3 are shown in Figure 7E. Similar results were confirmed for sucrose (Figures 7F-J). Thus, we detected no biological sex differences. Interestingly, our results agree with what the current SABV would have revealed (see Figure S2-3). Because only one group was identified for METH and sucrose (Figure 7), we proceeded with multivariate analysis. For every subject (data shown in Table S1) for METH, and for sucrose (data shown in Table S2-3), we conducted multivariate analysis to determine relatedness, or the lack thereof, of the 10 variables obtained (Figure 8–9). For METH and sucrose, R2H-shift rate was directly related to R2H-shift price (Figure 7F, 7L) – similar results were obtained for the relationship between R2H-shift price/rate and other variables. We decided to report only R2H-shift rate statistics.

**Figure 7.**
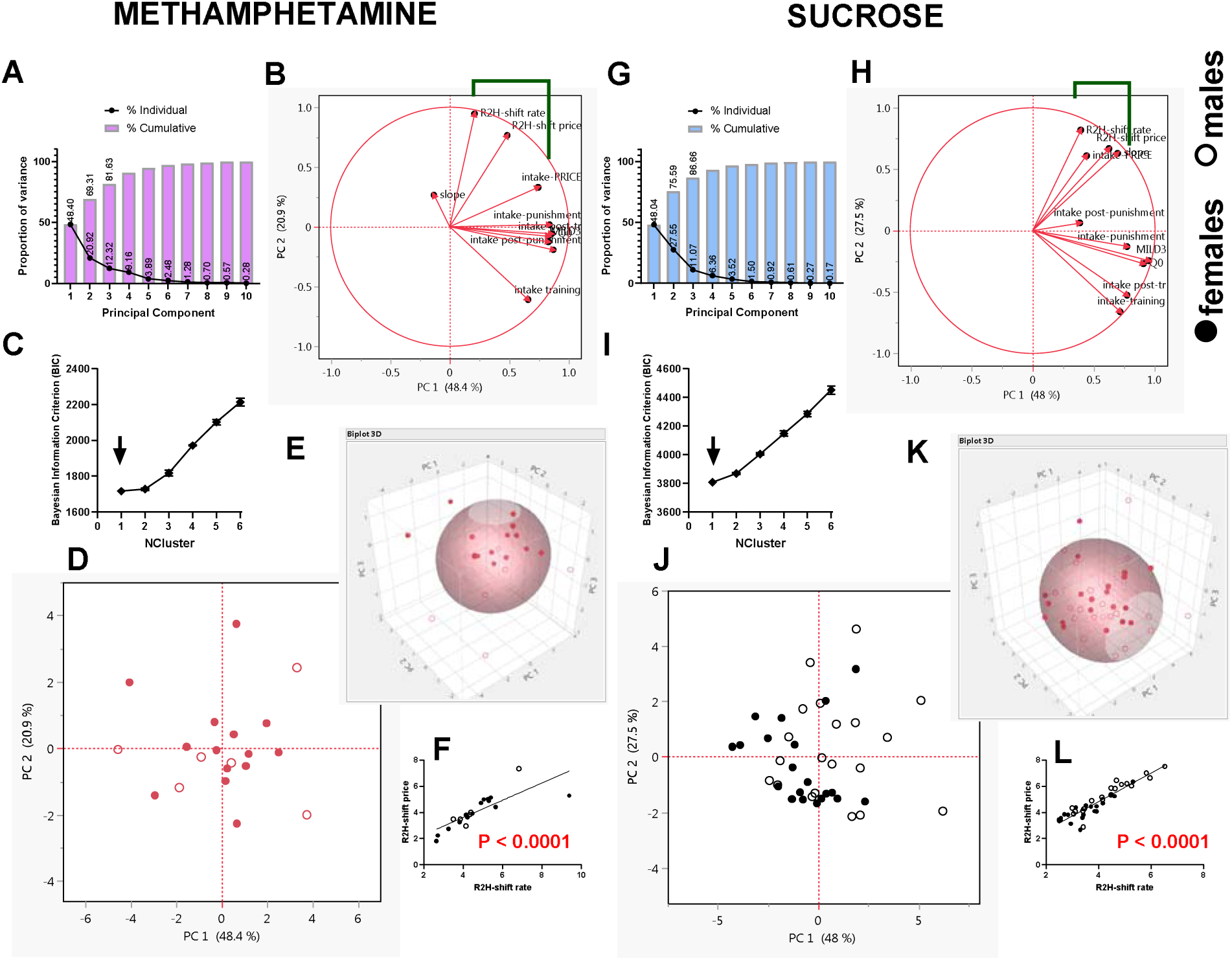
Clustering principal components revealed that all subjects for METH and sucrose studies belonged to one cluster: We conducted principal component analysis to combine the 10 variables obtained to derive principal components (PCs) for METH (A-F) and sucrose (G-L). A and G represent the principal components and their proportion of variance for METH and sucrose, respectively. B and H represent the variable loading for METH and sucrose, respectively. C and I reveal that the optimum number of clusters in the data set = 1 for METH and sucrose, respectively. D-E are 2-D and 3-D representation of the cluster identified for METH while J-K are the same for sucrose. In D-E and I-J open and closed circles represent males and females, respectively. F and L represent plots of the relationship between R2H-shift (x-axis) and R2H-shift price (y-axis) for METH and sucrose, respectively – these BEAST-derived R2H-shift variables were related (P < 0.0001).

**Figure 8:**
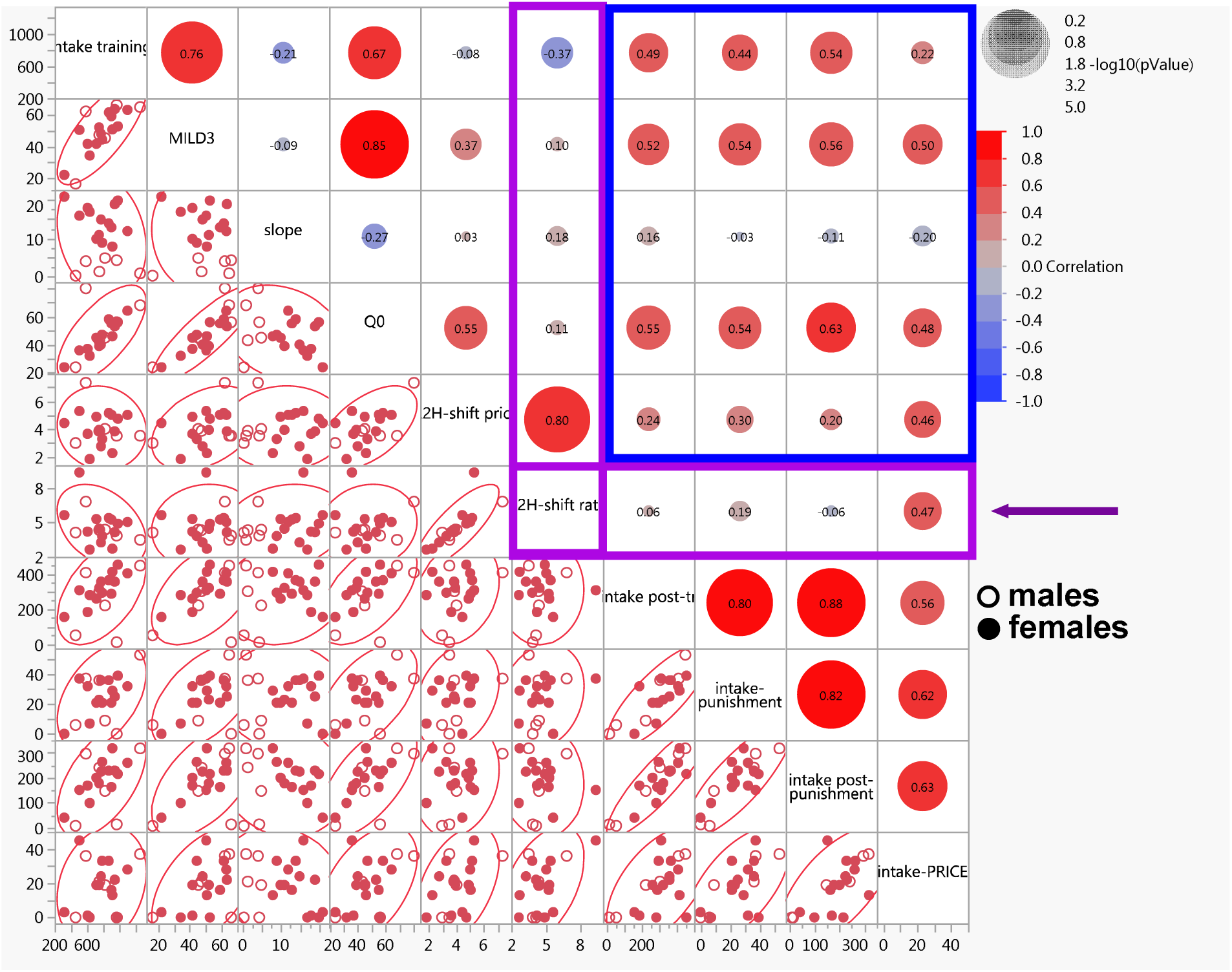
Multivariate analysis (METH study): R2H-shift rate is a novel variable and is unrelated to drug intake 1) during the training phase, and subsequent drug intake under 2) normal, 3) punishment, 4) post-punishment but related to intake under consumer-price constrained conditions. We conducted multivariate analysis for all 10 variables. The correlation coefficients are shown in the graph. The P values of the relationship(s) are shown as size of significance circles (the larger the circle, the stronger the relationship). Note that R2H-shift rate was related to intake-PRICE (area under the consumption-price response curve) (P < 0.05) but was unrelated to intake in the (1) post-training phase, 2) punishment phase, and 3) post-punishment phase (see horizontal arrow). Importantly, R2H-shift rate is truly novel – it was unrelated to any variables obtained concurrently during the self-administration training phase.

**Figure 9:**
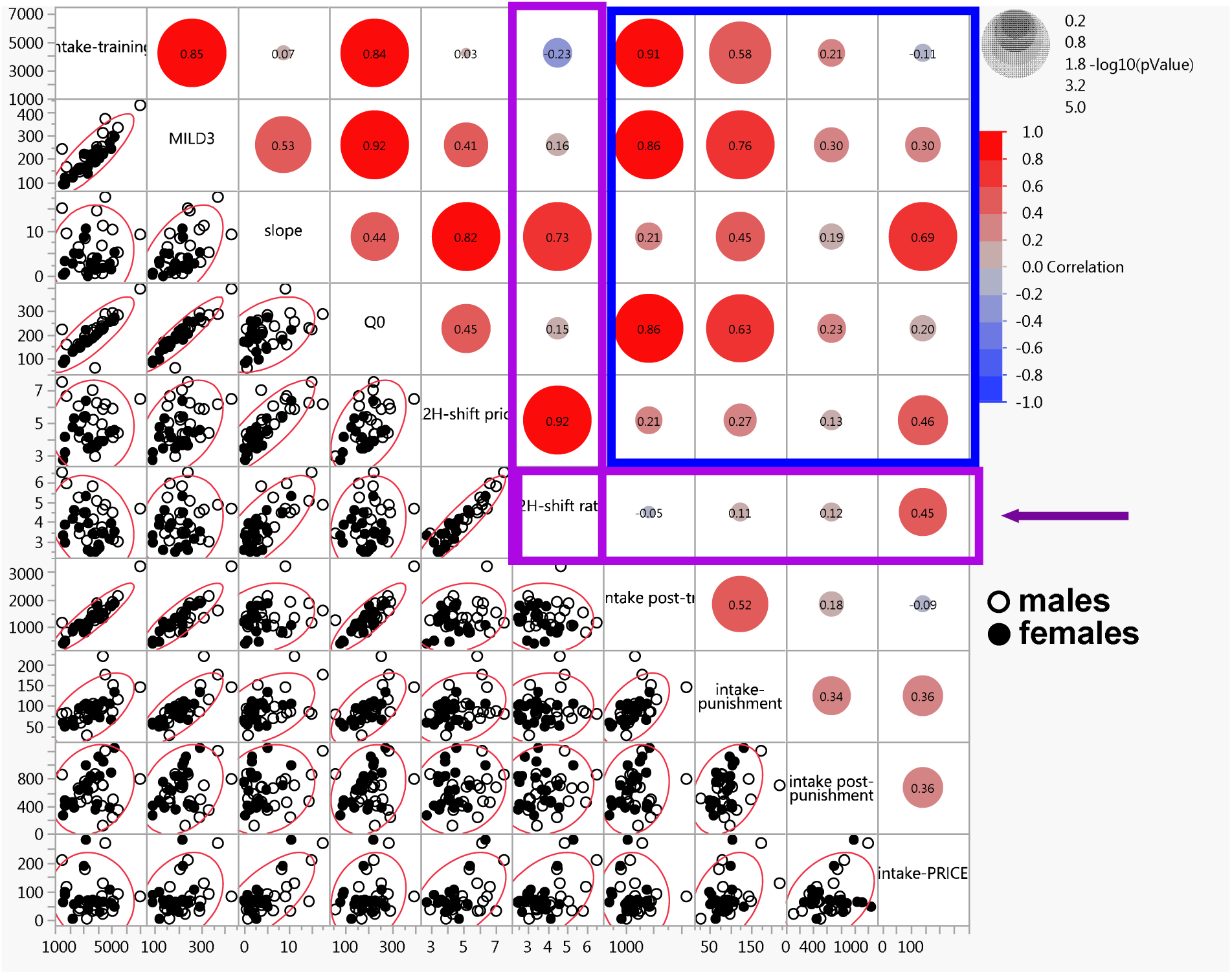
Multivariate analysis (sucrose study): R2H-shift rate is a novel variable and is unrelated to drug intake 1) during the training phase, and subsequent drug intake under 2) normal, 3) punishment, 4) post-punishment but related to intake under consumer-price constrained conditions. We conducted multivariate analysis for all 10 variables. The correlation coefficients are shown in the graph. The P values of the relationship(s) are shown as size of significance circles (the larger the circle, the stronger the relationship). Note that R2H-shift rate was related to intake-PRICE (area under the consumption-price response curve) (P < 0.05) but was unrelated to intake in the (1) post-training phase, 2) punishment phase, and 3) post-punishment phase (see horizontal arrow). Importantly, R2H-shift rate is truly novel – apart from slope, it was unrelated to any variables obtained concurrently during the self-administration training phase.

### Employment of Multivariate analysis to determine relationships between variables for METH and sucrose studies

#### METH study: Relationship between variables obtained within the training phase

R2H-shift rate was unrelated to intake-training (F 1, 18 = 2.930, P = 0.1041), MILD3 (F 1, 18 = 0.1766, P = 0.6793), slope (F 1, 18 = 0.5937, P = 0.4510) and Q_0_ (F 1, 18 = 0.2148, P = 0.6486). Q_0_ was related to intake-training (F 1, 18 = 14.73, P = 0.0012), MILD3 (F 1, 18 = 47.99, P < 0.0001), but not slope (F 1, 18 = 1.376, P = 0.2560). Other relationships are shown in Figure 8.

#### METH study: Relationship between training phase and post-training drug intake variables

For the METH study (Figure 8), intake-training could predict intake post-training (F 1, 18 = 5.696, P = 0.0282), intake-punishment (F 1, 18 = 4.423, P = 0.0498), intake post-punishment (F 1, 18 = 7.221, P = 0.0151), but not intake-PRICE (F 1, 18 = 0.9498, P = 0.3427). MILD3 could predict intake post-training (F 1, 18 = 6.590, P = 0.0194), intake-punishment (F 1, 18 = 7.254, P = 0.0149), intake post-punishment (F 1, 18 = 8.318, P = 0.0099) and intake-PRICE (F 1, 18 = 5.912, P = 0.0257). Slope could not predict any subsequent intake variables: intake post-training (F 1, 18 = 0.4879, P = 0.4938), intake-punishment (F 1, 18 = 0.01573, P = 0.9016), intake post-punishment (F 1, 18 = 0.2229, P = 0.6425) and intake-PRICE (F 1, 18 = 0.7295, P = 0.4043). BEAST derived parameter Q_0_ could predict all post-training drug intake variables - intake post-training (F 1, 18 = 8.007, P = 0.0111), intake-punishment (F 1, 18 = 7.317, P = 0.0145), intake post-punishment (F 1, 18 = 11.89, P = 0.0029) and intake-PRICE (F 1, 18 = 5.393, P = 0.0321). Intriguingly, R2H-shift rate could only predict intake-PRICE (F 1, 18 = 5.173, P = 0.0354). R2H-shift rate was unrelated to the other intake variables: intake post-training (F 1, 18 = 0.05918, P = 0.8106), intake-punishment (F 1, 18 = 0.6864, P = 0.4182) and intake post-punishment (F 1, 18 = 0.05905, P = 0.8108). Other relationships are shown in Figure 8.

#### Sucrose study: Relationship between variables obtained within the training phase

R2H-shift rate was related slope (F 1, 38 = 43.65, P < 0.0001), but not to intake-training (F 1, 38 = 2.100, P = 0.1555), MILD3 (F 1, 38 = 0.9676, P = 0.3315) and Q_0_ (F 1, 38 = 0.5774, P = 0.4520). Q_0_ was related to intake-training (F 1, 38 = 88.07, P < 0.0001), MILD3 (F 1, 38 = 203.8, P < 0.0001) and slope (F 1, 38 = 9.303, P = 0.0042). Other relationships are shown in Figure 9.

#### Sucrose study: Relationship between training phase and post-training drug intake variables

For the sucrose study (Figure 9), intake-training could predict intake post-training (F 1, 38 = 176.9, P < 0.0001) and intake-punishment (F 1, 38 = 19.74, P < 0.0001), but not intake post-punishment (F 1, 38 = 1.747, P = 0.1942) and not intake-PRICE (F 1, 38 = 0.4374, P = 0.5124). MILD3 could predict intake post-training (F 1, 38 = 107.1, P < 0.0001) and intake-punishment (F 1, 38 = 52.88, P < 0.0001), but neither intake post-punishment (F 1, 38 = 3.694, P = 0.0621) nor intake-PRICE (F 1, 38 = 3.841, P = 0.0574). Slope could predict intake punishment (F 1, 38 = 9.801, P = 0.0033) and intake-PRICE (F 1, 38 = 33.62, P < 0.0001) but could predict neither intake post-training (F 1, 38 = 1.793, P = 0.1885) nor intake post-punishment (F 1, 38 = 1.469, P = 0.2330). BEAST derived variable Q_0_ (like intake-training) could predict intake post-training (F 1, 38 = 56.64, P < 0.0001) and intake-punishment (F 1, 38 = 15.78, P = 0.0003), but not intake post-punishment (F 1, 38 = 1.498, P = 0.2285) and not intake-PRICE (F 1, 38 = 1.796, P = 0.1881). Intriguingly, R2H-shift rate could only predict intake-PRICE (F 1, 38 = 9.399, P = 0.0040), but not intake post-training (F 1, 38 = 0.09066, P = 0.7650), intake-punishment (F 1, 38 = 0.4442, P = 0.5091) and intake post-punishment (F 1, 38 = 0.5239, P = 0.4736). Other relationships are shown in Figure 9.

The summary of our results are shown in Figure 10.

**Figure 10.**
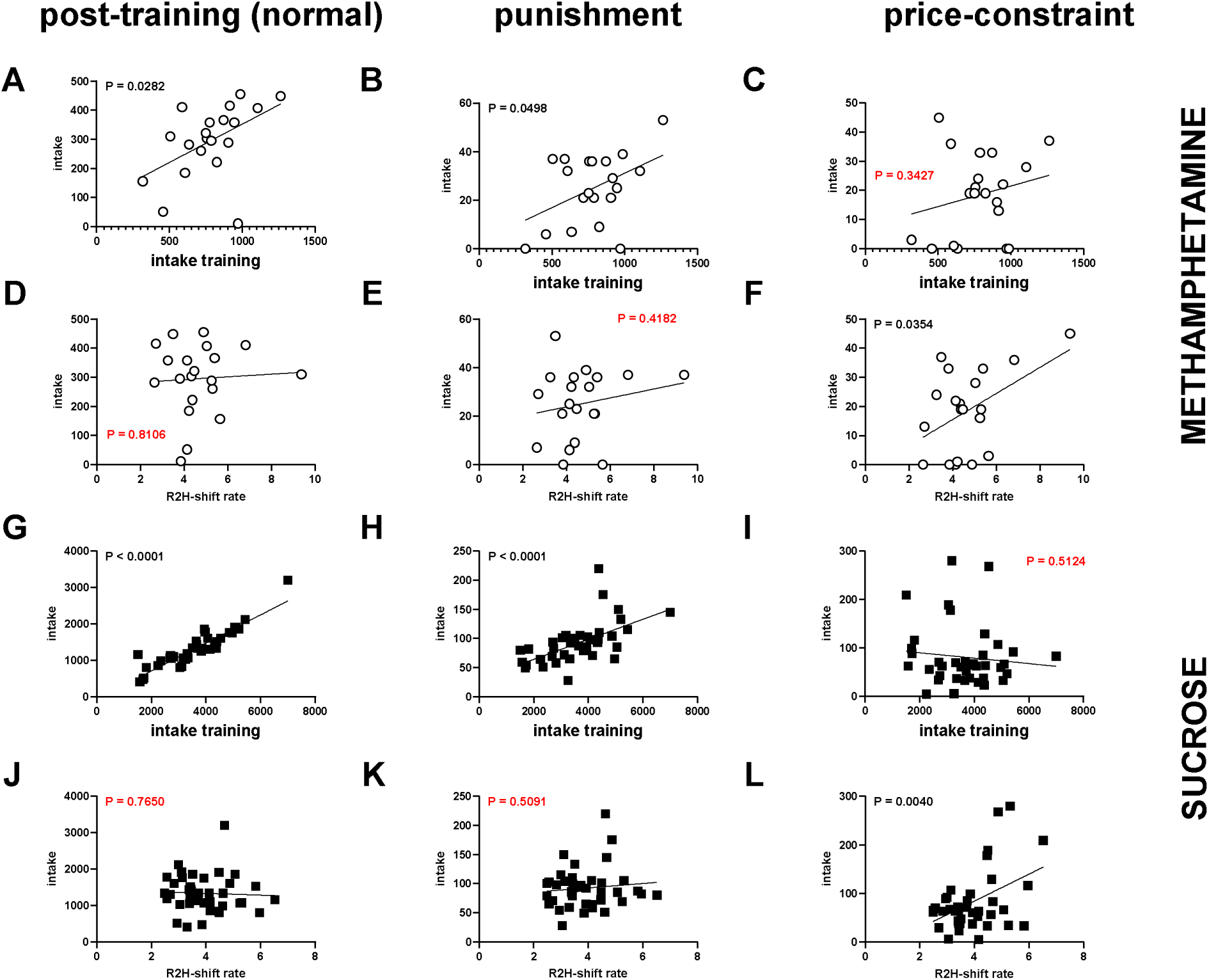
For METH and sucrose, R2H-shift rate is related to economic demand, but is unrelated to drug intake levels (under normal and punishment conditions): For METH (A-F) and sucrose (G-L), we compared/contrasted drug intake levels and R2H-shift rate obtained during the training phase for effectiveness in predicting drug intake levels after the training phase under normal (left column), punishment (middle column) and price constrained (right column) conditions. Graphs A-C and G-I are for relationships between prior intake (intake training phase) and intake under normal/punishment and price constrained conditions for METH and sucrose. Graphs D-F and J-L are for relationships between R2H-shift rate (obtained during the training phase) and intake under normal/punishment and price constrained conditions for METH and sucrose. Linear regression analysis revealed that intake training could predict subsequent intake under normal/punishment conditions for METH (A-B) and sucrose (G-H). Interestingly, intake training could not predict intake under the consumption-price curve for METH (C) and sucrose (I). Conversely, linear regression analysis revealed that R2H-shift rate could not predict subsequent intake under normal/punishment conditions for METH (D-E) and sucrose (J-K). Interestingly, R2H-shift rate could predict intake under the consumption-price curve for METH (F) and sucrose (L). In summary, R2H-shift rate is related to economic demand, but is unrelated to drug intake levels (under normal and punishment conditions)

## Discussion

With the idea that the shift from recreational to habitual drug use is governed by behavioral economic principles, we developed a novel model (BEAST) to quantify R2H-shift during drug self-administration time course (Figure 1). Our goal was to determine if R2H-shift variables during self-administration training phase was/were related to 1) drug intake during the self-administration time course, and subsequent drug intake under 2) normal, 3) punishment, 4) post-punishment, and 5) price-constrained conditions. Our experimental protocol, shown in Figure 2, was designed to capture several variables per subject during and after the R2H-shift. We obtained 10 parameters including variables from current and BEAST for every individual – male and female (Figure 3–6). Because data from the saline study could not be fitted by BEAST (Figure S1), this group was excluded from further analysis. Using PCA/ gaussian mixtures model clustering of PCs derived from combination of all 10 variables, we determined that there were no biological sex differences for METH and sucrose (Figure 7). Not only were there no biological sex differences, but there were also no distinct groups (Figure 7). Because there was only one group identified, we conducted multivariate analysis on all data (Figure 8–9). With regards to our goals, we determined that R2H-shift variables were unrelated to drug intake during the self-administration time course, and during subsequent drug intake under normal/punishment/post-punishment conditions (Figure 8–10). Interestingly, R2H-shift variables were related (only) to drug intake during price-constrained conditions (Figure 10). Because R2H-shift may represent a motivation-habitual construct, our results reinforce the idea that intake and motivation are not necessarily related (Castaneda and Job, 2026), see also (Bentzley et al., 2014, 2013; Ikemoto and Panksepp, 1996; Oleson et al., 2011; Roberts et al., 2013; Siciliano and Jones, 2017).

There are some limitations of this study. One limitation include the low n of males employed in the METH study (n = 6). That said, we started with adequate n (16), but BEAST revealed that 10 of these male rats were not motivated for the drug. The low number of males would have been a limitation for SABV, but we have shown that SABV is an extremely limited approach for understanding drug use/abuse (Job, 2024; Showell and Job, 2024; Tigano and Job, 2025, 2024; Yao and Job, 2024).

In line with DSM-V definitions that person(s) that have developed substance use disorders use the substances despite negative consequences, a preclinical punishment model has been developed and is assumed to be effective in separating drug users into non-compulsive (or shock-sensitive, SS) and compulsive (or shock-resistant, SR) phenotypes. SS and SR METH users are thought to represent non-compulsive and compulsive drug users, respectively (Cadet et al., 2019, 2017, 2016; Campbell et al., 2018; Daiwile et al., 2024; Datta et al., 2018b, 2018a; Domi et al., 2021; Duan et al., 2022; Durand et al., 2021; Giuliano et al., 2021, 2019, 2018; Hopf and Lesscher, 2014; Hu et al., 2019; Jayanthi et al., 2022a, 2022b; Krasnova et al., 2017; Marchant et al., 2018; Munoz et al., 2023; Pelloux et al., 2015, 2012, 2007; Subu et al., 2020; Sun and Yuill, 2020; Torres et al., 2018, 2017; Xue et al., 2012; Zhou et al., 2019). Since R2H-shift may represent the transition from non-compulsive to compulsive, SR R2H-shift should be greater than SS R2H-shift, but this was not the case (Figure S4). Moreover, grouping subjects as SS versus SR groups (see Figure S4) may not represent true groups as our cluster-based model (including of novel BEAST variables) revealed that there was only one cluster (Figure 7). Prior METH experience may be an important factor predicting SS/SR (Figure 8). Indeed, this is supported by evidence suggesting that the response of METH consumption to neuropharmacological manipulation is related to prior METH experience (Chojnacki et al., 2020; Culbertson et al., 2009; Kazahaya et al., 1989; Lorrain et al., 2000; Mcfadden et al., 2013; McFadden et al., 2015, 2012; Orio et al., 2010; Segal et al., 2003; Vezina et al., 1999; Wee et al., 2007) and footshock stress may be conceptualized as a type of neuropharmacological manipulation – it alters the release of several endogenous mediators, including dopamine (Dazzi et al., 2004, 2001a, 2001b; Kalivas and Duffy, 1995; Motzo et al., 1996; Puglisi-Allegra et al., 1991; Saulskaya et al., 2008; Sorg and Kalivas, 1993, 1991; Takahashi et al., 1998; Yoshioka et al., 1996; Young, 2004; Young and Rees, 1998). If SS/SR are more similar than they are different, the punishment model may not be an effective tool for behavioral phenotyping for substance use disorders, though more work needs to be done to clarify this.

Current drug user typology separates drug takers into high and low takers (HT versus LT) based solely on their drug intake under normal conditions. We showed that intake was unrelated to R2H-shift (Figure 8–9) and HT versus LT do not have distinct R2H-shift variables (Figure S5). Thus, HT versus LT groups (see Figure S5) may not represent true groups as our cluster-based model (including novel BEAST variables) revealed that there was only one cluster (Figure 7). Distinguishing subjects based on drug intake alone does not appear to be effective for behavioral phenotyping for substance use disorders, we have already established this (Castaneda and Job, 2026), though more work needs to be done to clarify this.

In summary, we developed/validated a behavioral economic model which allows us to assess the shift from recreational-to-habitual drug use, and this may open up opportunities to understand this mechanism. In this study, we report that the recreational-to-habitual shift associated with the progression of drug use predicted/was related to economic demand (a measure of motivation) but not drug intake levels (under normal and punishment conditions).

## Acknowledgements

The authors acknowledge the support of NIH grant **DA054461** (MOJ). The content is solely the responsibility of the authors and does not necessarily represent the official views of the National Institutes of Health. Additional support was provided by Rowan University via the Francis R. Lax Fund for Faculty Development (MOJ), and faculty startup funds (MOJ).

## SUPPLEMENTAL FILES

## Supplemental Table Legends

**Table S1:** Variables obtained for the male (M) and female (F) subjects that self-administered methamphetamine (METH) and self-administration time demand curves could be fit using the exponential model. Note that n = 10 males were excluded and n = 0 females were excluded.

| RatID | SEX | Intake training | MILD3 | slope | Q<sub>0</sub> | R2H-shift price | R2H-shift rate | Intake post-training | Intake-punishment | Intake post-punishment | Intake PRICE |
| --- | --- | --- | --- | --- | --- | --- | --- | --- | --- | --- | --- |
| 1 | M | 458 | 16.67 | 0.32 | 24 | 2.963 | 4.135 | 52 | 6 | 8 | 0 |
| 5 | M | 970 | 66 | 4.32 | 56 | 3.487 | 3.853 | 11 | 0 | 14 | 0 |
| 6 | M | 756 | 47.33 | 1.35 | 43 | 3.846 | 4.341 | 303 | 36 | 241 | 21 |
| 7 | M | 587 | 62 | 4.02 | 80 | 7.341 | 6.823 | 410 | 37 | 296 | 36 |
| 8 | M | 1264 | 64.67 | 0.87 | 68 | 3.488 | 3.493 | 449 | 53 | 317 | 37 |
| 14 | M | 825 | 45.33 | 4.87 | 45 | 3.995 | 4.388 | 222 | 9 | 147 | 19 |
| 17 | F | 916 | 50.67 | 7.87 | 46 | 2.242 | 2.708 | 415 | 29 | 318 | 13 |
| 18 | F | 788 | 44.33 | 8.87 | 47 | 3.261 | 3.812 | 295 | 21 | 263 | 33 |
| 19 | F | 717 | 41.67 | 9.87 | 45 | 4.876 | 5.306 | 261 | 21 | 200 | 19 |
| 20 | F | 751 | 52 | 10.87 | 39 | 3.919 | 4.467 | 322 | 23 | 178 | 19 |
| 21 | F | 1105 | 63 | 11.87 | 64 | 4.999 | 5.042 | 407 | 32 | 260 | 28 |
| 22 | F | 905 | 59 | 12.87 | 55 | 5.023 | 5.252 | 288 | 21 | 163 | 16 |
| 23 | F | 872 | 61.33 | 13.87 | 58 | 5.137 | 5.404 | 366 | 36 | 228 | 33 |
| 24 | F | 775 | 48 | 14.87 | 40 | 2.745 | 3.256 | 358 | 36 | 224 | 24 |
| 25 | F | 505 | 50.33 | 15.87 | 36 | 5.284 | 9.382 | 310 | 37 | 151 | 45 |
| 26 | F | 635 | 34.33 | 16.87 | 32 | 1.810 | 2.639 | 282 | 7 | 99 | 0 |
| 27 | F | 607 | 41.67 | 17.87 | 37 | 3.633 | 4.219 | 185 | 32 | 169 | 1 |
| 28 | F | 946 | 63.33 | 18.87 | 53 | 3.819 | 4.143 | 358 | 25 | 229 | 22 |
| 29 | F | 986 | 52.67 | 19.87 | 56 | 4.725 | 4.899 | 455 | 39 | 215 | 0 |
| 30 | F | 317 | 22.33 | 20.87 | 24 | 4.421 | 5.645 | 156 | 0 | 40 | 3 |

**Table S2:** Variables obtained for male subjects that self-administered sucrose and self-administration time demand curves could be fit using the exponential model. Note that n = 2 males were excluded.

| RatID | SEX | Intake training | MILD3 | slope | Q<sub>0</sub> | R2H-shift price | R2H-shift rate | Intake post-training | Intake-punishment | Intake post-punishment | Intake PRICE |
| --- | --- | --- | --- | --- | --- | --- | --- | --- | --- | --- | --- |
| 31 | M | 5102 | 295.67 | 2.99 | 260 | 4.083 | 3.109 | 1915 | 150 | 494 | 67 |
| 32 | M | 7000 | 428.67 | 9.13 | 394 | 6.453 | 4.676 | 3203 | 145 | 793 | 83 |
| 34 | M | 3646 | 216.67 | 3.73 | 219 | 7.016 | 5.820 | 1525 | 87 | 672 | 33 |
| 35 | M | 3699 | 209.33 | 3.92 | 209 | 5.180 | 4.123 | 1336 | 105 | 678 | 52 |
| 36 | M | 3827 | 187.00 | 0.56 | 58 | 2.888 | 3.414 | 1253 | 79 | 867 | 38 |
| 37 | M | 2816 | 159.00 | 3.86 | 157 | 4.973 | 4.153 | 1099 | 58 | 458 | 63 |
| 38 | M | 3949 | 250.00 | 14.40 | 228 | 6.210 | 5.075 | 1857 | 85 | 470 | 66 |
| 39 | M | 5057 | 291.00 | 10.22 | 292 | 5.860 | 4.477 | 1910 | 85 | 237 | 33 |
| 40 | M | 4339 | 219.33 | 0.49 | 224 | 4.290 | 3.466 | 1389 | 93 | 705 | 38 |
| 41 | M | 4364 | 230.00 | 2.17 | 227 | 4.278 | 3.440 | 1453 | 99 | 112 | 23 |
| 42 | M | 5433 | 332.00 | 5.12 | 282 | 4.343 | 2.996 | 2122 | 115 | 343 | 92 |
| 44 | M | 2733 | 144.00 | 1.99 | 146 | 4.115 | 3.403 | 1048 | 83 | 574 | 43 |
| 45 | M | 3255 | 162.00 | 0.78 | 170 | 3.886 | 3.045 | 1025 | 28 | 507 | 6 |
| 46 | M | 2680 | 153.67 | 6.40 | 160 | 6.028 | 5.247 | 1062 | 69 | 660 | 34 |
| 47 | M | 1507 | 241.33 | 14.98 | 222 | 7.521 | 6.520 | 1162 | 80 | 856 | 209 |
| 48 | M | 4381 | 310.33 | 11.15 | 254 | 5.837 | 4.628 | 1336 | 220 | 700 | 129 |
| 49 | M | 3120 | 208.33 | 8.31 | 191 | 5.336 | 4.465 | 835 | 72 | 757 | 178 |
| 50 | M | 1811 | 163.67 | 9.43 | 158 | 6.638 | 5.957 | 803 | 82 | 453 | 116 |
| 51 | M | 4534 | 370.00 | 17.48 | 286 | 6.145 | 4.871 | 1602 | 175 | 1204 | 268 |
| 52 | M | 4316 | 251.67 | 6.71 | 235 | 4.903 | 3.738 | 1340 | 97 | 401 | 85 |

**Table S3:**
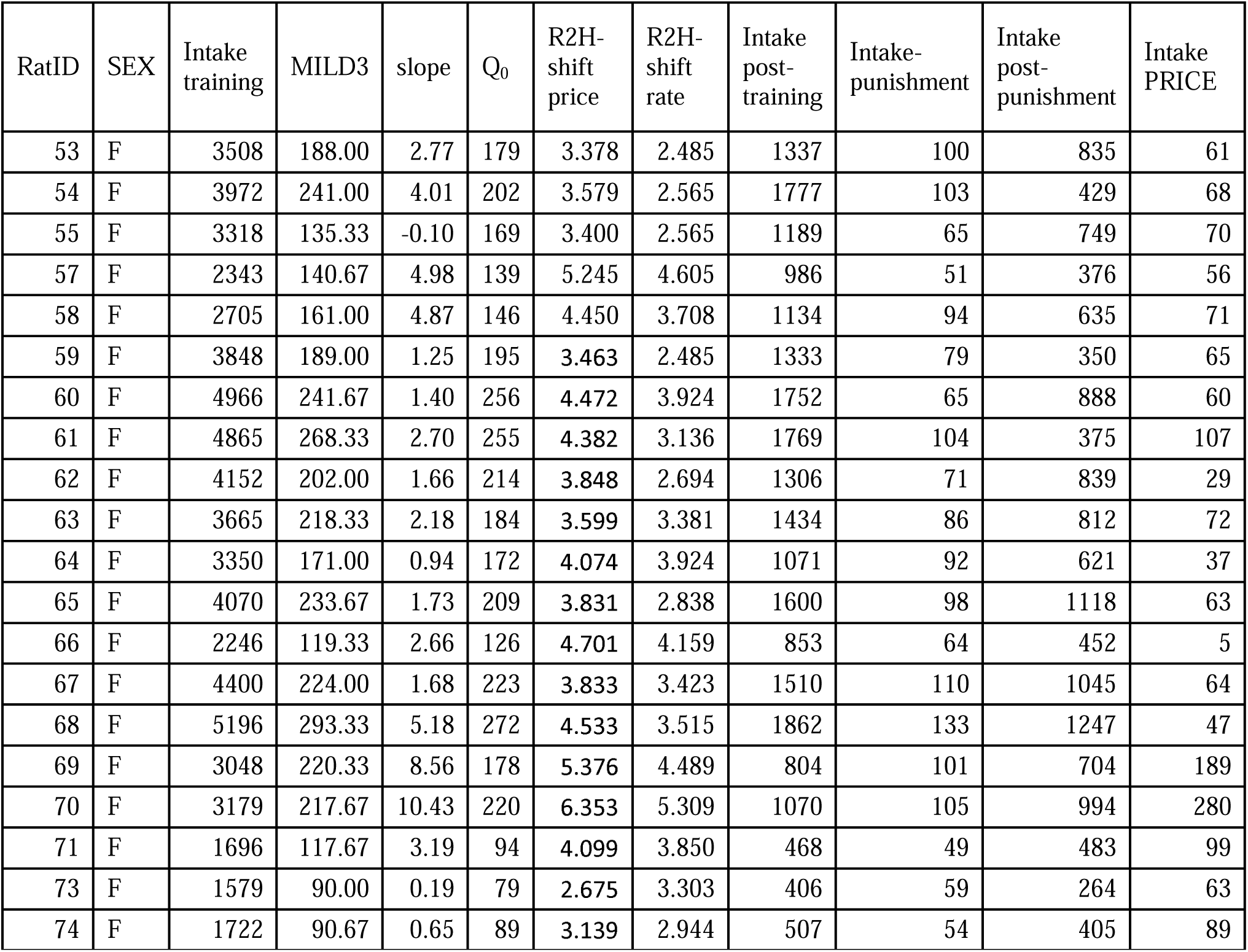
Variables obtained for female subjects that self-administered sucrose and self-administration time demand curves could be fit using the exponential model. Note that n = 2 females were excluded.

## Supplemental Figure Legends

**Figure S1.**
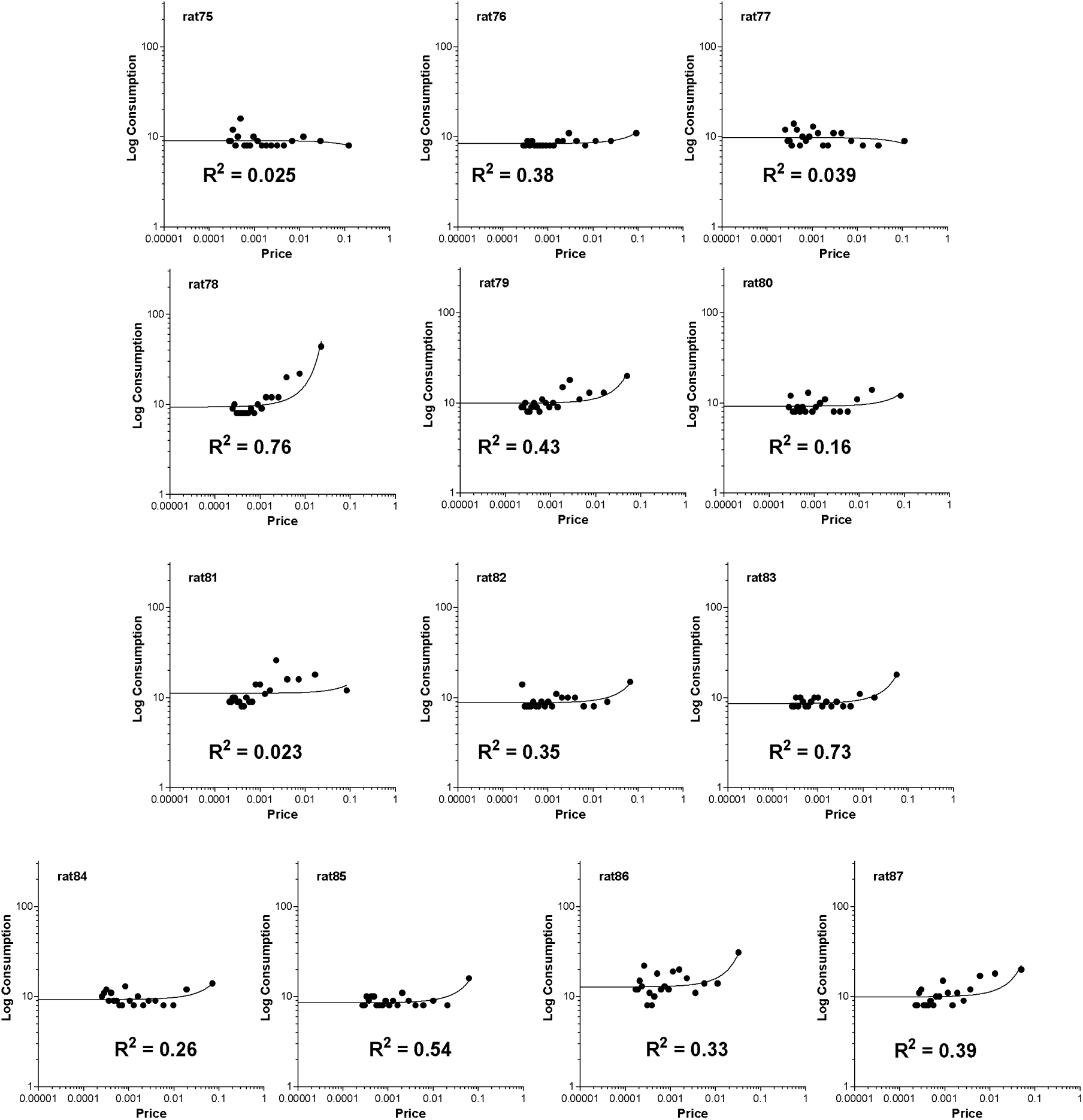
The demand curves of individuals (males and females) derived from saline self-administration time curve analysis using the new Behavioral Economic model for the Analysis of Self-administration Time-curve (BEAST). Each graph represents an individual. The n = 3 male rats that self-administered saline were labeled as rat75 – rat77 while the n = 10 female rats that self-administered saline were labeled as rat78-rat87. The R^2^ for goodness of fit is written in each graph. Note that the exponential model for demand curves could not fit and/or yielded negative values for eValue and Pmax for all subjects. These individuals (the entire saline group) were excluded from further analysis.

**Figure S2.**
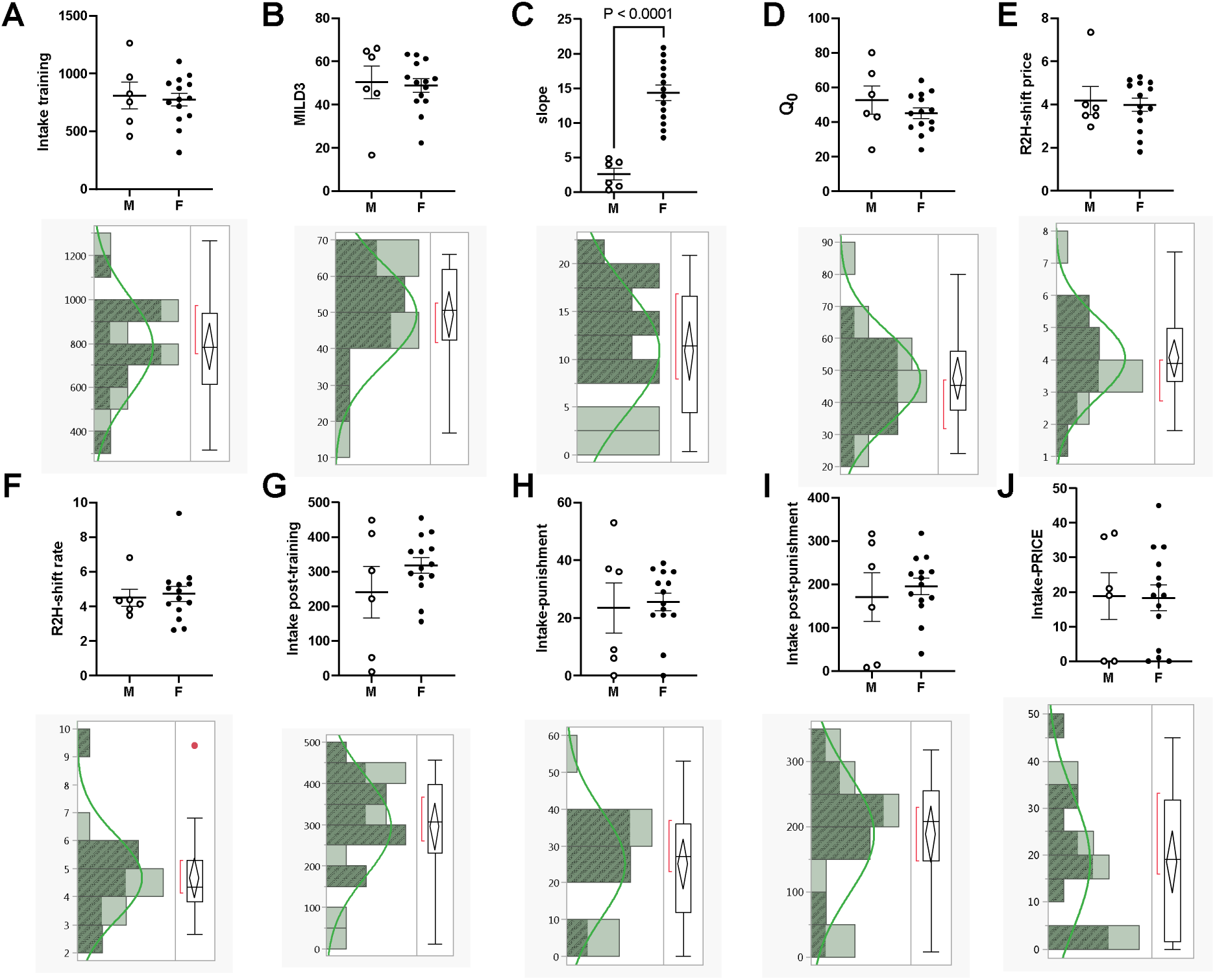
Analysis of the distribution for all of the 10 variables suggests that males and females self-administering METH are not distinct groups. The criteria for confirming distinct groups by biological sex is 1) there should be two clearly defined populations, and 2) these populations should mostly be non-overlapping. The graphs A-J show male versus female comparisons for all 10 variables. The histograms below the graphs A-J represent output from analysis of the distribution. The shaded and unshaded regions of the histograms represent females and males, respectively. The box and violin plots next to the histograms show the median value for the variables. There were significant differences between males and females only for slope (C) but not for all other variables (A-B, D-J). For all 10 variables, distribution analysis suggested the data represented a single normally-distributed population. By variable count, it may imply that males and females were different for 10% of the total number of variables analyzed but similar for 90% of the total number of variables analyzed. By percentage assessment males and females were more similar than they were different, overall.

**Figure S3.**
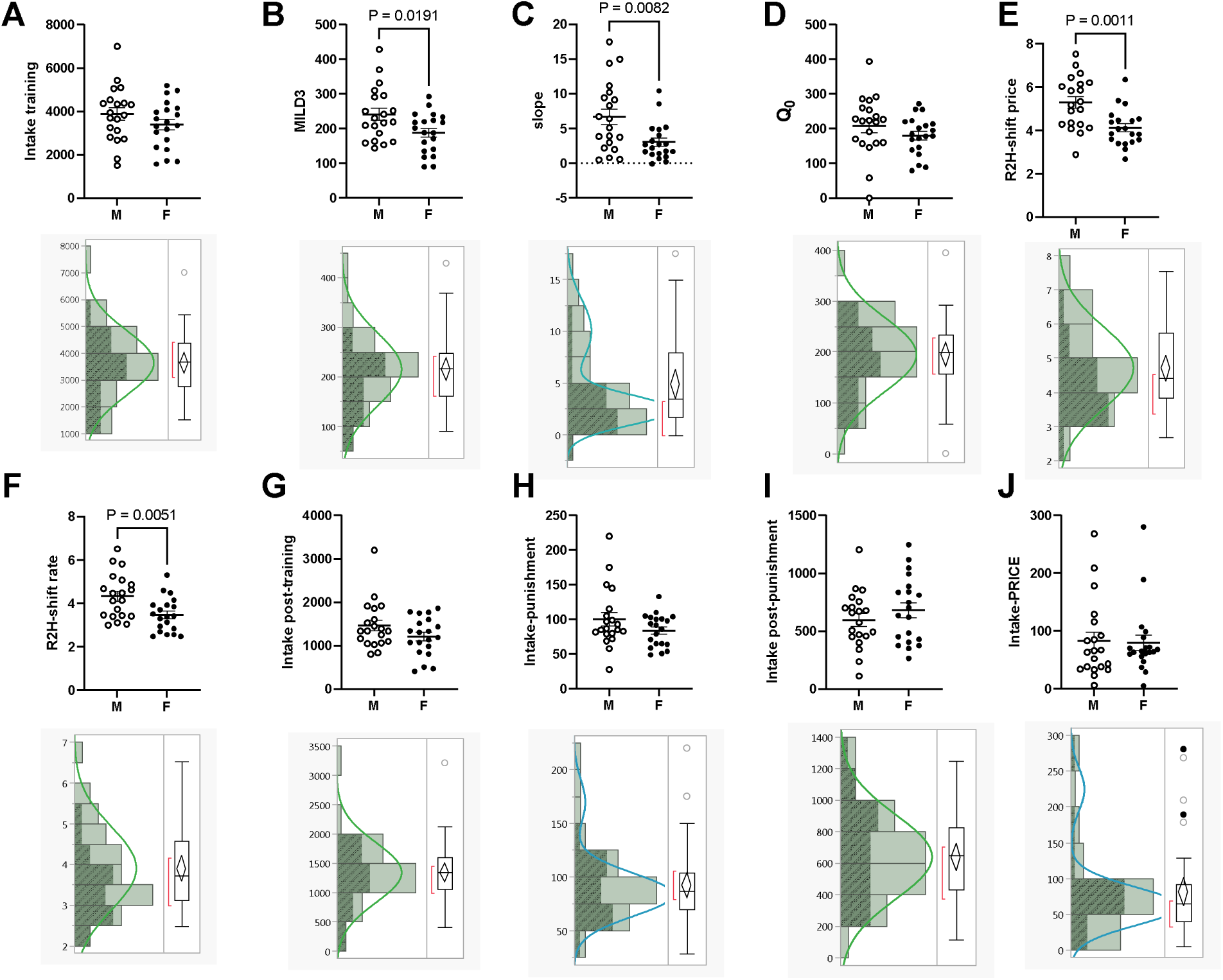
Analysis of the distribution for each of the 10 variables suggests that males and females self-administering sucrose are not distinct groups. The criteria for confirming distinct groups by biological sex is 1) there should be two clearly defined populations, and 2) these populations should mostly be non-overlapping. The graphs A-J show male versus female comparisons for all 10 variables. The histograms below the graphs A-J represent output from analysis of the distribution. The shaded and unshaded regions of the histograms represent females and males, respectively. The box and violin plots next to the histograms show the median value for the variables. There were significant differences between males and females for MILD3, slope, R2H-shift rate, R2H-shift price (B, C, E and F) but not for all other variables (A, D, G-J). While different, MILD3, R2H-shift price and R2H-shift rate represented a single normally-distributed population (B, E and F). But even for slope (C), which revealed a bimodal distribution, the populations were overlapping, not clearly distinct, with each comprised of both males and females. By variable count, it may imply that males and females were different for 40% of the total number of variables analyzed but similar for 60% of the total number of variables analyzed. By percentage assessment males and females were more similar than they were different, overall.

**Figure S4.**
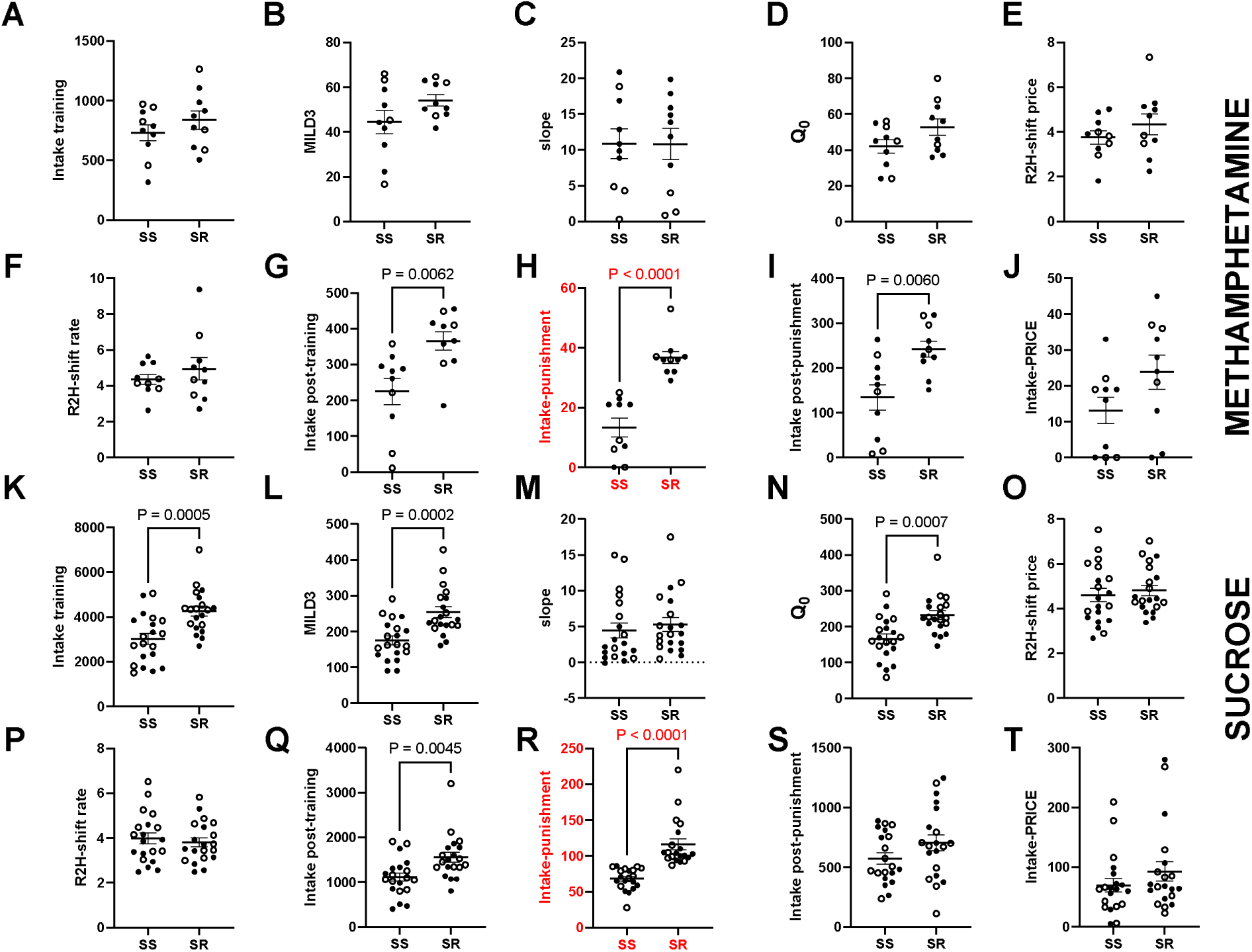
Punishment-resistant versus Punishment-sensitive do not represent groups with distinct recreational-to-habitual shift rates for the drug: Open and close circles represent males and females, respectively. With the rationale that punishment-resistant (or shock-resistant, SR) and punishment-sensitive (or shock-sensitive, SS) are similar to high versus low takers under punishment conditions, we employed median split of their intake under punishment to separate the groups (see red highlighted graphs in G for METH and Q for sucrose). The other graphs are comparison of other variables of the same designated SR versus SS takers. For METH, SR versus SS takers (H) were (also) distinct for intake post-training (G) and intake post-punishment (I) – they were not distinct for intake-PRICE (J) or other intake variables (A-D) and R2H-shift variables (E-F). For sucrose, SR versus SS were distinct for intake post-training (K) and, except for slope, were distinct for all prior intake-type variables (intake-training, MILD3, Q_0_, K,L and O). Note that sucrose SS and SR groups were not different for the R2H-shift price/rate (O- P). In summary, for METH and sucrose, SS and SR were distinct prior to the punishment phase (G, Q), but they were not different with regards to intake under the consumption-price curve (J, T) and they were not different with regards to R2H-shift variables (E-F, O-P).

**Figure S5.**
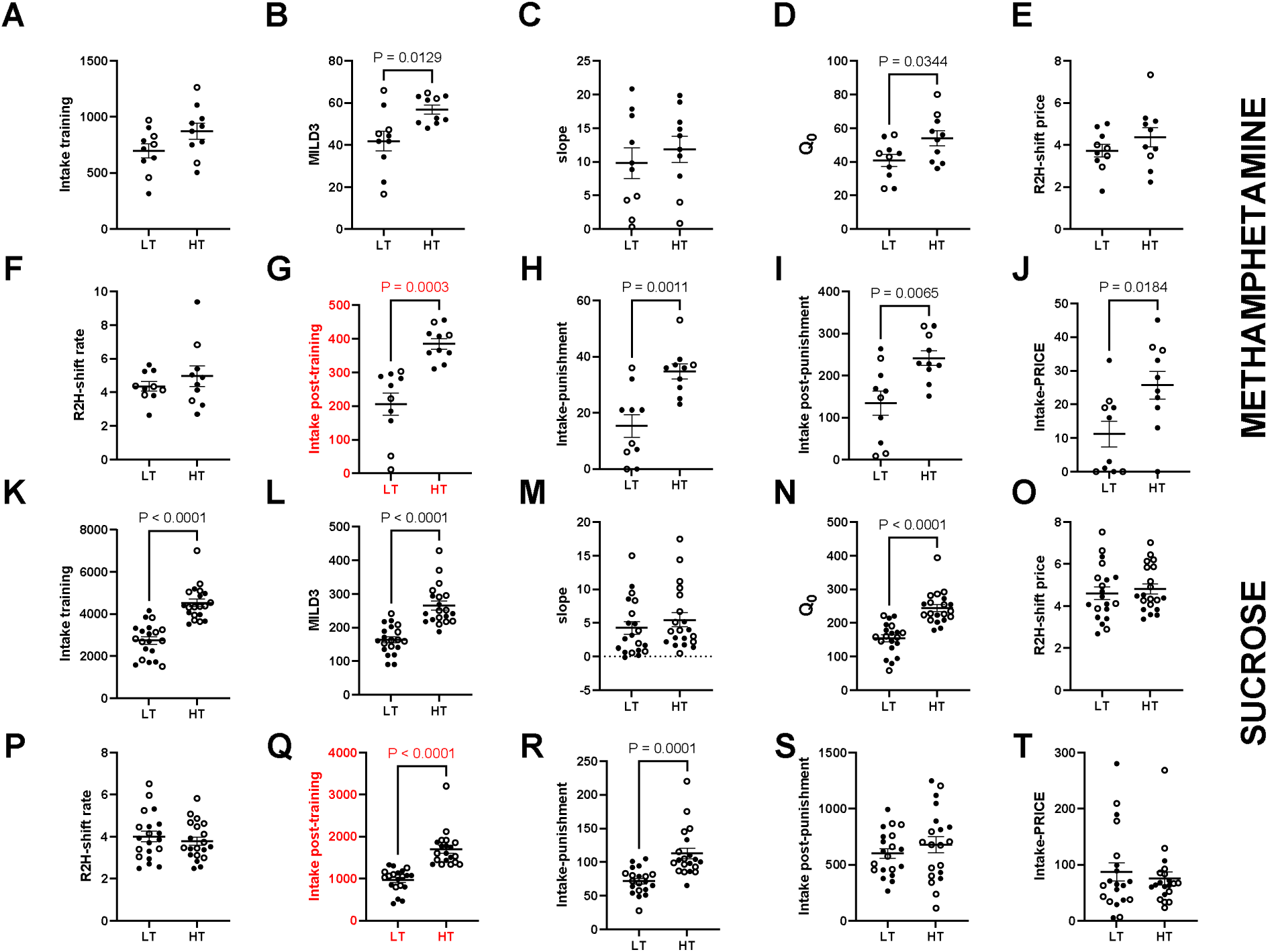
High versus Low Takers do not represent groups with distinct recreational-to-habitual shift rates for the drug: Open and close circles represent males and females, respectively. The graphs A-J and K-T, respectively, represent comparison between high and low takers (HT and LT) for 10 variables for METH and sucrose, respectively. High and low takers were derived using median split of intake post-training (see red highlighted graphs in G for METH and Q for sucrose). The other graphs are comparison of other variables of the same designated high versus low takers. For METH, HT and LT (G) were distinct for most intake variables obtained during the training phase (MILD3 and Q_0_, B and D), and all intake after the training phase (H-J). HT and LT were not distinct with respect to R2H-shift price/rate (E-F). For sucrose, HT and LT (Q) were distinct for most intake variables obtained during the training phase (intake training, MILD3 and Q_0_, A-B and D), and intake during the punishment phase (R). HT and LT were not distinct with respect to R2H-shift price/rate (O-P). In summary, for METH and sucrose, HT and LT were not distinct with regards to R2H-shift price/rate (E-F, O-P).

